# Structural tuning of native type V fimbriae shapes mechanical specialization in *Porphyromonas gingivalis*

**DOI:** 10.64898/2026.07.16.738973

**Authors:** Zhifeng Wang, Adwaith B Uday, Valentin Nelea, Dieter P. Reinhardt, D. Peter Tieleman, Khanh Huy Bui, Natalie Zeytuni

## Abstract

Fimbriae are proteinaceous filaments central to bacterial adhesion, colonization, and biofilm formation. *Porphyromonas gingivalis*, a keystone periodontal pathogen, utilizes two type V fimbriae systems, the major Fim and the minor Mfa fimbriae, that support distinct adhesive and biofilm-associated functions. Here, we determine the high-resolution cryo-electron microscopy structures of natively assembled FimA and Mfa1 stalks purified directly from *P. gingivalis.* Despite a shared donor-strand exchange mechanism, the two stalks exhibit distinct helical geometries and donor-strand environments. FimA has a larger helical pitch, whereas the Mfa1 donor strand is more extensively buried and further shielded by an ordered N-terminal latch. Comparison with monomeric stalk pilin structures reveals distinct assembly-associated remodeling around the donor-strand interface. Deletion of the Mfa1 latch reduces heat-resistant oligomer accumulation and produces species consistent with incomplete processing, yet recovered filaments retain the donor-strand-exchanged stalk architecture and exhibit initial force peaks comparable to native Mfa1 during simulated axial extension. Atomic force microscopy reveals greater apparent local stiffness of FimA under indentation, whereas steered molecular dynamics simulations show multiple lower-force axial transitions in FimA but a dominant higher-force transition in Mfa1. In both systems, the exchanged donor strands remain engaged during simulated extension while distinct surrounding contact networks rearrange. Together, these findings demonstrate how a conserved polymerization linkage can support structurally and mechanically distinct adhesive filaments, providing a molecular basis for mechanical diversification of type V fimbriae.

## Introduction

*Porphyromonas gingivalis* is a keystone oral pathogen that contributes to the development and progression of periodontal disease and has been associated with systemic inflammatory and degenerative conditions, including cardiovascular disease, rheumatoid arthritis, diabetes, and Alzheimer’s disease^1–3^. Its ability to colonize and persist within subgingival polymicrobial communities relies on interactions with host tissues and neighboring microorganisms ^4–7^. Among the principal surface structures supporting these interactions are two antigenically and functionally distinct type V fimbriae systems, traditionally described as the long, major Fim fimbriae and the shorter, minor Mfa fimbriae ^8–10^. Fim fimbriae participate in broad adhesive interactions with host cells, extracellular matrix components, salivary proteins, and microbial partners^11–15^, whereas Mfa fimbriae have prominent roles in interbacterial interactions, including coadhesion with early colonizers such as *Streptococcus gordonii*, autoaggregation, microcolony formation, and biofilm maturation^9,16–18^. These distinct but partly overlapping functions position Fim and Mfa as important mediators of both host attachment and integration into microbial communities.

Both systems share a modular organization comprising a polymeric stalk assembled through C-terminal donor-strand exchange, tip-associated accessory proteins, and a membrane-anchored base that contributes to surface retention and filament length regulation (**Fig. 1A**)^10^. These modules are formed by distinct sets of proteins: FimA forms the Fim stalk, FimC-E are the accessory or tip-associated components, and FimB provides the anchoring and length-regulatory functions. The corresponding components of the Mfa system are Mfa1, Mfa3-5, and Mfa2, respectively^10^. Perturbation of FimB or Mfa2 alters filament length and surface retention, illustrating that filament organization is controlled at multiple levels^19,20^.

**Figure 1.**
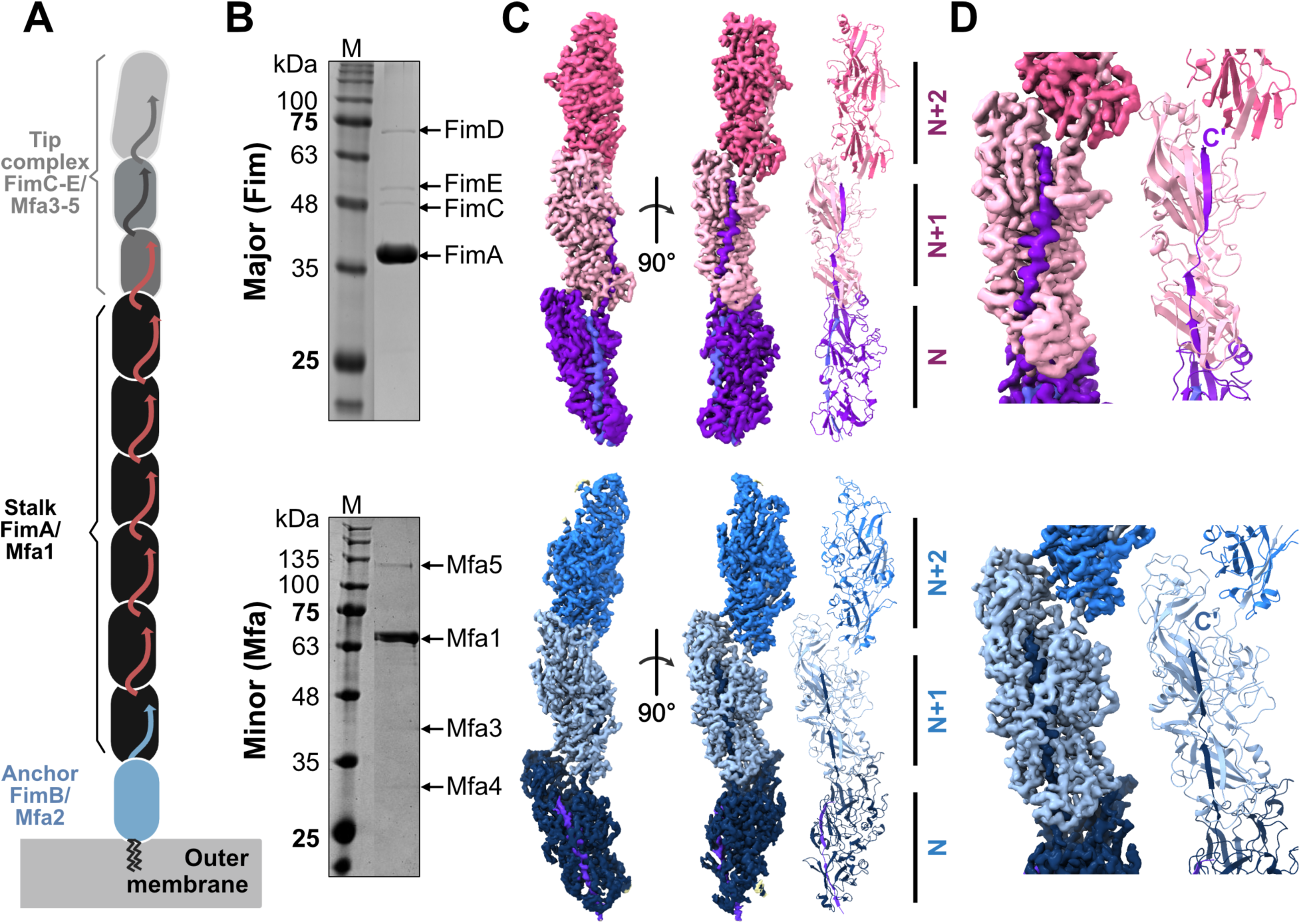
Cryo-EM structure of native Fim and Mfa fimbriae stalks. **(A)** Schematic organization of the major Fim and minor Mfa fimbriae of *P. gingivalis*. The fimbriae consist of an outer-membrane anchor (FimB or Mfa2), a polymeric stalk formed by FimA or Mfa1, and distal tip components comprising FimC-FimE or Mfa3-Mfa5, respectively. Red arrows indicate the C-terminal donor strands that extend from one stalk pilin subunit into the neighboring subunit during donor-strand exchange. **(B)** SDS-PAGE analysis of purified Fim and Mfa fimbriae. Expected masses of mature stalk pilins and tip components: FimA (36.5 kDa), FimC (43.9 kDa), FimD (70.0 kDa), FimE (55.2 kDa), Mfa1 (55.6 kDa), Mfa3 (45.0 kDa), Mfa4 (31.0 kDa), and Mfa5 (112.0 kDa). Mfa1 and Mfa5 migrate at anomalously high apparent molecular weights on SDS-PAGE^10,30^. M: protein marker. **(C)** Cryo-EM reconstructions and fitted atomic models of the FimA (top) and Mfa1 (bottom) polymerized stalks, shown across three successive stalk pilin subunits. Subunits are labeled N, N+1, and N+2 and colored by their successive positions along the polymer. The FimA stalk adopts a more extended helical architecture, with a pitch of 340.2 Å, a rise of 68.2 Å and a twist of 72.1°, whereas the Mfa1 stalk forms a more compact filament, with a pitch of 279.1Å, a rise of 71.7 Å and a twist of 92.4°. **(D)** Close-up views of the donor-strand exchange interfaces in FimA and Mfa1 stalk pilins. In both stalks, the C-terminal donor strand from one stalk pilin subunit occupies the acceptor groove of the adjacent subunit. In Mfa, the exchanged donor strand is more deeply buried within the acceptor groove and is further shielded by an ordered N-terminal region, producing a more enclosed donor-strand interface than in Fim.

The Fim and Mfa systems are encoded by distinct operons, with the overall organization of each operon broadly conserved across *P. gingivalis* strains^15^. Within this conserved organization, both stalk pilins exhibit substantial sequence diversity. FimA variants are classified into types I-V, whereas Mfa1 comprises the 53 and 70 groups, with the latter further divided into 70A and 70B^15^. These variants differ in adhesive or antigenic properties, indicating that sequence diversification occurs within otherwise conserved fimbriae systems^21,22^.

Despite this diversity, FimA and Mfa1 share a conserved two-domain β-sandwich architecture and are produced as lipoprotein precursors^15,23^. Following export to the cell surface, these precursors undergo N-terminal processing by arginine-specific gingipains (RgpA or RgpB)^24^. Proteolytic maturation enables conversion from intramolecular self-complementation to intermolecular donor-strand exchange, in which the C-terminal donor strand of one stalk pilin inserts into the acceptor groove of a neighboring subunit and completes its β-sheet architecture^25^. Unlike classical chaperone-usher assembly, this pathway does not require a separate chaperone to supply a complementing strand. Previous studies have shown that recombinant FimA and Mfa1 expressed in *Escherichia coli* can polymerize *in vitro* following RgpB-mediated cleavage^10,24,26^. Although RgpA/B gingipains mediate canonical N-terminal maturation, residual Mfa1 polymerization has been observed in the absence of canonical Rgp cleavage, suggesting that alternative proteolytic pathways can also contribute to maturation^24,27^.

High-resolution structures of recombinant monomeric pilins and polymerized stalks have established the molecular basis of protease-mediated type V fimbriae assembly^10,15,24^. However, stalk organization is relevant to more than polymer formation. In both Fim and Mfa fimbriae, the adhesive interactions are not restricted to the tip-associated accessory components, as the stalk pilins themselves have been implicated in binding host molecules and microbial partners. The polymeric stalk therefore combines adhesive and structural roles, making its molecular architecture relevant both to the presentation of interaction sites and to the forces transmitted through them.

Studies of other bacterial pili demonstrate that stalk deformation can regulate force transmission to adhesive contacts. For example, reversible uncoiling of *Escherichia coli* type 1 fimbriae modulates the load transmitted to the terminal adhesin^28^. For Fim and Mfa, the shared donor-strand exchange mechanism explains how adjacent pilins remain connected but does not establish how the stalks accommodate deformation. Axial extension could involve displacement of the exchanged donor strand, rearrangement of neighboring contacts, or conformational changes within the pilins themselves. Distinguishing among these possibilities is necessary to understand how a conserved assembly mechanism can support different mechanical responses.

Here, we combine high-resolution cryo-electron microscopy (cryo-EM), atomic force microscopy (AFM), steered molecular dynamics (SMD), and structure-guided perturbation to examine the structural and mechanical specialization of native FimA type I and Mfa1-70A stalks. Filaments purified directly from *P. gingivalis* reveal distinct helical geometries, donor-strand environments, and assembly-associated remodeling. AFM and SMD identify distinct responses to local indentation and axial extension, while the simulations show that deformation is accommodated by rearrangement of surrounding contact networks without loss of donor-strand engagement. Together, these findings reveal how a conserved donor-strand linkage can support distinct deformation pathways, providing a molecular basis for mechanical diversification in bacterial adhesive polymers.

## Results

### Native FimA and Mfa1 fimbriae adopt distinct helical architectures

To define the architectures of the two principal type V fimbriae systems of *P. gingivalis*, we purified natively assembled Fim (nFim) and Mfa (nMfa) fimbriae from ATCC 33277 Δ*mfa1::erm^R^* and ATCC 33277 Δ*fimA::erm^R^* strains, respectively (**Fig. 1B**). Under the purification and imaging conditions used, nFim fimbriae were recovered as long filaments, frequently extending beyond 2 µm, whereas nMfa fimbriae were predominantly shorter, typically in the ∼100 nm range (**Fig. S1A**). These observations are consistent with previous reports of short wild-type Mfa fimbriae^15,24^ and the elongated Fim phenotype of ATCC 33277^19^. The latter is associated with an intrinsic mutation in *fimB*, which encodes the anchoring and length-regulatory protein FimB^19^.

Single-particle cryo-EM analysis yielded reconstructions of both stalks at 2.9 Å resolution, enabling manual tracing and assignment of atomic models comprising three consecutive stalk pilin subunits for each filament (**Fig. 1C, Fig. S1, Table S1**). The first resolved residue was Ala47 in FimA, consistent with the reported N-terminal maturation sites associated with arginine-specific gingipain processing^15,25^. In Mfa1, the mature terminus is Ala50^27^, but this residue was unresolved. Our model begins at Gly51, likely due to the local flexibility. The two filaments formed right-handed helical assemblies with distinct geometries. The nFimA stalk adopted a more extended architecture with a pitch of 340.2 Å, an axial rise of 68.2 Å and a twist of 72.1° whereas nMfa1 stalk adopted a more compact architecture, with a pitch of 279.1 Å, an axial rise of 71.7 Å and a twist of 92.4° (**Fig. S2A, B)**. Despite these differences in helical packing, the two stalks share the characteristic type V donor-strand exchange mechanism. The C-terminal donor strand of each stalk pilin inserts into the acceptor groove of the adjacent subunit, completing its β-sheet architecture. The exchanged strands are stabilized by backbone hydrogen bonding and extensive hydrophobic interactions with residues lining the acceptor groove (**Fig. 1D; Fig. S2C**). The local organization of this shared interface nevertheless differs markedly between FimA and Mfa1. In nMfa1, the exchanged donor strand is more deeply buried within the acceptor groove than the counterpart in nFimA (**Fig. 1D**). An ordered N-terminal region (residues 51-64) further encloses the Mfa1 interface by folding over the exchanged donor strand, forming a latch-like element without an equivalent covering arrangement in nFimA (**Fig. 1D**). We used Protein interfaces, surfaces and assemblies’ service PISA^29^ to analyze and quantify the burial of the C-terminal donor strand at the inter-subunit interface. In FimA, the donor strand had a calculated solvent-accessible surface area (ASA) of 3113.4 Å^2^, of which 2221.8 Å^2^ was buried through inter-subunit association, corresponding to 71.4%. In Mfa1, 2698.6 Å^2^ of the corresponding donor-strand ASA of 3,147.0 Å^2^ was buried, corresponding to 85.8%. The buried fraction was therefore 14.4% higher in Mfa1. These measurements demonstrate a greater enclosure of the Mfa1 donor strand and establish differences in solvent exposure around a shared inter-subunit linkage.

Two additional non-protein densities were observed in the nMfa1 map. The first lies within a loop containing Asp507, Asp509, Asn512, Glu514, and Asn515, corresponding to the proline-rich metal-binding region previously observed in recombinant Mfa1 (rMfa1) structures (**Fig. S3**)^24^. Although calcium was not added during purification, the density was assigned as Ca²⁺ based on its coordination environment and agreement with the previously characterized site^24^. A second density was observed adjacent to the buried Arg483, at a position modeled as acetate in the recombinant monomeric Mfa1 crystal structure (rMfa1_monomer_; PDB 5NF2^10^) based on the contents of the crystallization conditions. The density dimensions are consistent with a small molecule or ion (**Fig. S3**). Since acetate was not included in our purification protocol, we also examined whether a covalent modification could account for this feature. Intact-protein mass spectrometry identified a dominant species at 55,645.56 Da, only 1.74 Da below its theoretical mass of 55,647.30 Da (**Fig. S3B**), with no additional resolved species corresponding to arginine monomethylation, dimethylation, acetylation, or phosphorylation^31^. Thus, the identity of the Arg483-associated density could not be assigned unambiguously, and no ligand was modeled at this site.

Finally, we compared the native polymerized stalk pilin conformations with those in previously reported recombinant assemblies generated by *in vitro* polymerization of proteins expressed in *E. coli*. Despite the different assembly and purification contexts, the corresponding subunit conformations showed close agreement. Superposition of nFimA with recombinant polymerized FimA (rFimA_polymer_; PDB 6KMF^25^) yielded a Cα RMSD of 0.74 Å across 337 aligned residue pairs. The nMfa1 and recombinant polymerized Mfa1 (rMfa1_polymer_; PDB 21CO^24^) superimposed with a Cα RMSD of 0.91 Å across 513 aligned residue pairs. These comparisons establish close correspondence between the polymerized pilin conformations in recombinant assemblies and those in filaments assembled by *P. gingivalis*.

### Assembly-associated conformational remodeling differentially reorganizes the donor-strand environment in FimA and Mfa1

To identify assembly-associated conformational differences, we compared the native polymerized stalk subunits with previously reported monomeric structures of recombinant rFimA_monomer_ (PDB 6JZK^25^) and rMfa1_monomer_ (PDB 5NF2^10^). These comparisons reveal local rearrangements around the acceptor groove between the intramolecularly complemented monomeric conformations and the intermolecularly donor-strand-exchanged conformations in the assembled stalks (**Fig. 2A, B; Movies S1, S2**).

**Figure 2.**
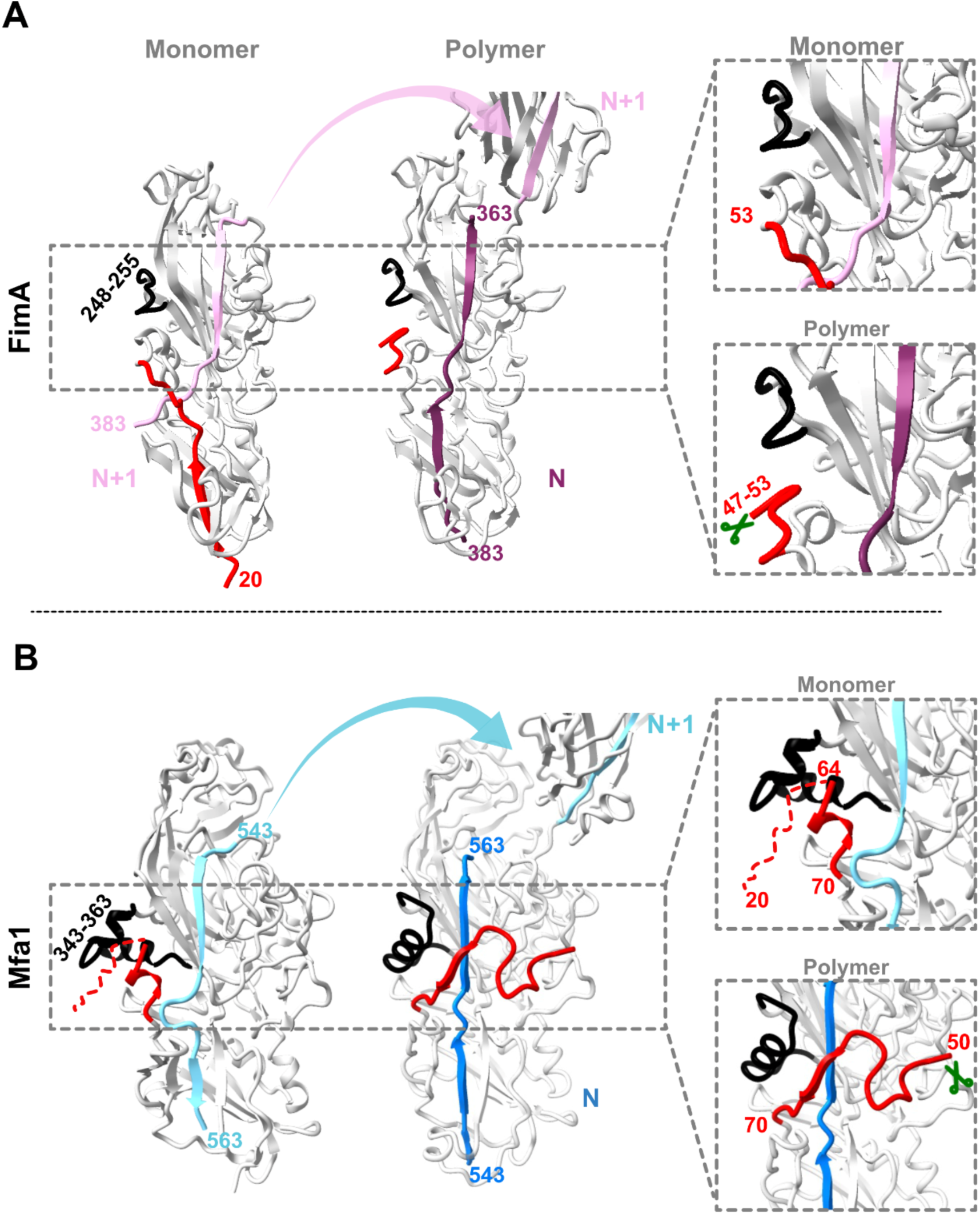
Donor-strand exchange is accompanied by distinct assembly-associated conformational remodeling in FimA and Mfa1. **(A)** Comparison of monomeric pre-polymerized FimA (PDB ID 6JZK^25^) and polymerized FimA within the native Fim stalk. FimA maturation occurs by N-terminal proteolytic processing by arginine-specific gingipain proteases, indicated by green scissors, with Ala47 forming the mature N-terminal residue in the polymerized stalk. This comparison highlights donor-strand exchange and assembly-associated conformational remodeling surrounding the acceptor groove. In the polymerized state, the C-terminal donor strand from subunit N+1 inserts into the acceptor groove of subunit N, replacing the intramolecular self-complementing strand observed in the monomeric state. Regions that undergo local conformational remodeling include the N-terminal region (residues 47-53) and an adjacent loop-like segment (residues 248-255), shown in red and black, respectively. **(B)** Comparison of monomeric pre-polymerized Mfa1 (PDB 5NF2^10^) and polymerized Mfa1 within the native Mfa stalk. Mfa1 maturation occurs by N-terminal proteolytic processing before polymer incorporation; the first resolved residue in the native Mfa stalk model is Gly51. This comparison highlights donor-strand exchange and assembly-associated conformational remodeling surrounding the acceptor groove. In addition to donor-strand exchange, polymerization remodels two donor-strand-adjacent elements: the Mfa1 N-terminal region (residues 51-70) in red, which folds over the exchanged donor strand, and an adjacent α-helical segment (residues 343-363) in black. The red dashed line indicates the N-terminal latch (51-64) that is disordered in the monomeric Mfa1 crystal structure but becomes ordered in the polymerized stalk.

In FimA, assembly-associated remodeling involves the retained N-terminal segment spanning residues 47-53 and the adjacent loop spanning residues 248-255 (**Fig. 2A**). The N-terminal segment lies near the self-complementing donor strand in the monomeric structure but adopts a different conformation in the assembled stalk, where it does not form a covering element over the exchanged donor strand. Repositioning of the neighboring 248-255 loop further alters the local organization beside the donor-strand interface.

In Mfa1, residues 51-64, which are not represented in the monomeric model, form an ordered N-terminal latch over the exchanged donor strand supplied by a neighboring subunit (**Fig. 2B**). The adjacent α5 region of the polymerized pilin (α5_Polymer_), spanning residues 343-363, also changes conformation and position relative to the monomeric state. The FimA 248-255 loop and Mfa1’s α5 occupy structurally analogous positions beside the donor-strand interface, identifying corresponding sites of assembly-associated remodeling despite their different secondary structures.

We next examined variation in these remodeling elements across representative FimA genotypes I-V and the Mfa1-70A, Mfa1-70B, and Mfa1-53 groups, using sequence alignments and comparisons of the native reference structures with AlphaFold3-predicted models (**Fig. 3**). The predicted FimA structures retained the overall pilin fold, whereas the donor-strand-adjacent loop corresponding to ATCC 33277 residues 248-255 varied in sequence and length. Relative to the ATCC 33277 FimA-I reference, this segment showed no sequence similarity in some genotypes and up to 87.5% similarity in others. It occupied a similar position in the predicted structures across the five genotypes but was shorter in genotypes IV and V (**Fig. 3A**).

**Figure 3.**
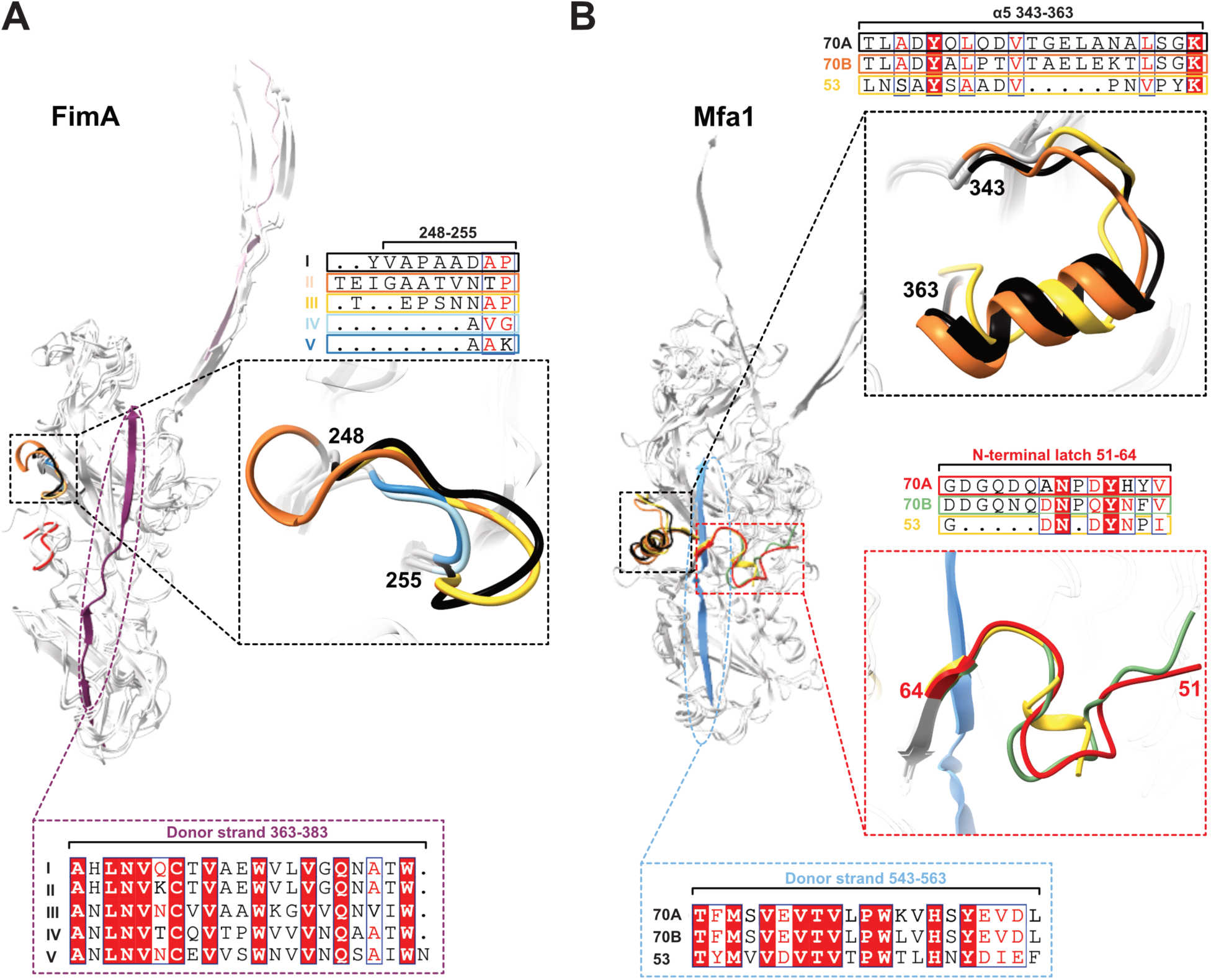
Structural conservation of assembly-associated remodeling elements across FimA and Mfa1 variants. **(A)** Sequence and structural comparison of representative FimA genotypes I-V. The structure-guided alignment shows the donor-strand-adjacent region corresponding to residues 248-255 of FimA-I from *P. gingivalis* ATCC 33277. The structural superposition shows the corresponding loop, with genotypes I, II, III, IV, and V in black, orange, yellow, light blue, and dark blue, respectively; the remainder of the structure is shown predominantly in light gray. The exchanged C-terminal donor-strand region corresponding to residues 363-383 of ATCC 33277 FimA-I is shown at the bottom. Residue numbering refers to the ATCC 33277 FimA-I reference sequence. (**B**) Sequence and structural comparison of representative Mfa1-70A, Mfa1-70B, and Mfa1-53 variants. Structure-guided alignments show the α5 helix region spanning residues 343-363, the N-terminal latch (51-64), and the exchanged C-terminal donor strand spanning residues 543-563 of Mfa1-70A from ATCC 33277. Structural superpositions show the α5 helix region (70A, black; 70B, orange; 53, yellow) and the N-terminal latch region (70A, red; 70B, green; 53, yellow). Residue numbering refers to Mfa1-70A from ATCC 33277. Conserved and similar residues are highlighted according to ESPript conventions. Structure-guided alignments were generated using FoldMason^33^ and visualized with ESPript 3.0^34,35^. The experimentally determined ATCC 33277 FimA-I and Mfa1-70A structures were used as references, whereas structures representing the remaining variants were predicted using AlphaFold3 in polymerized, donor-strand-exchanged configurations.

The Mfa1 α5 region likewise showed substantial sequence variation. Relative to ATCC 33277 Mfa1-70A, the corresponding segment in Mfa1-70B showed 66.7% similarity, whereas Mfa1-53 showed 28.6% sequence similarity and contained a five-residue deletion in the alignment. An α-helical element was nevertheless predicted at the corresponding position in each group. The Mfa1-70A and Mfa1-70B models showed similar local organization, while the predicted Mfa1-53 helix was shorter (**Fig. 3B**). Thus, preservation of an α5-like element did not require uniform conservation of its sequence or helical extent.

Variation also occurred in the N-terminal region corresponding to the Mfa1 latch (residues 51-64). Across the regions corresponding to ATCC 33277 Mfa1-70A latch, Mfa1-70B and Mfa1-53 showed 78.6% and 42.9% sequence similarity, respectively, with substitutions and alignment gaps particularly evident in Mfa1-53 (**Fig. 3B**). Additional differences occurred in the upper loops of the Mfa1 fold, where Mfa1-53 diverged from the 70A and 70B groups in both sequence and predicted local conformation. This sequence divergence is compatible with filament formation: previous studies demonstrated assembly following replacement of Mfa1-70 with Mfa1-53 and directly visualized fimbriae purified from the Mfa1-53 strain Ando^22,32^.

In contrast to several of these surrounding remodeling elements, the exchanged C-terminal donor strands were comparatively well conserved. Relative to the corresponding ATCC 33277 references, the FimA donor-strand region spanning residues 363-383 showed 61.9-100% sequence similarity across genotypes I-V, while the Mfa1 region spanning residues 543-563 showed 81.0-100% similarity across the 70A, 70B, and 53 groups (**Fig. 3A, B**).

Together, the native structures identify assembly-associated repositioning and distinct N-terminal arrangements around the donor-strand connection. The genotype comparisons place several of these remodeling elements within regions of variable sequence and predicted local organization, flanking comparatively conserved donor strands. This distinction provided the basis for examining how the surrounding contacts accommodate mechanical loading and for elucidating the contribution of the Mfa1 latch to polymer formation and assembled-stalk properties.

### Steered molecular dynamics reveals distinct tensile response regimes in FimA and Mfa1 polymerized stalks

Having identified differences in super-helical architecture, assembly-associated conformations, and donor-strand enclosure, we next examined how FimA and Mfa1 stalks respond to axial tensile loading. We performed SMD simulations using two-subunit stalk fragments derived from the nFimA and nMfa1 cryo-EM determined structures. Each fragment included an additional standalone donor-strand segment positioned in the acceptor groove of one terminal subunit to maintain strand complementation at that end of the model. A defined region of one subunit was restrained, while the distal terminal region of the neighboring subunit was pulled along the longitudinal axis of the fragment (**Fig. 4A**). Simulations were conducted at five pulling velocities spanning 0.02-0.50 nm ns⁻¹ under matched conditions to compare force-extension profiles and assess the pulling-rate dependence of structural transitions (**Figs. S4 and S5**). Force values were interpreted comparatively within this protocol rather than as estimates of physiological rupture forces

**Figure 4.**
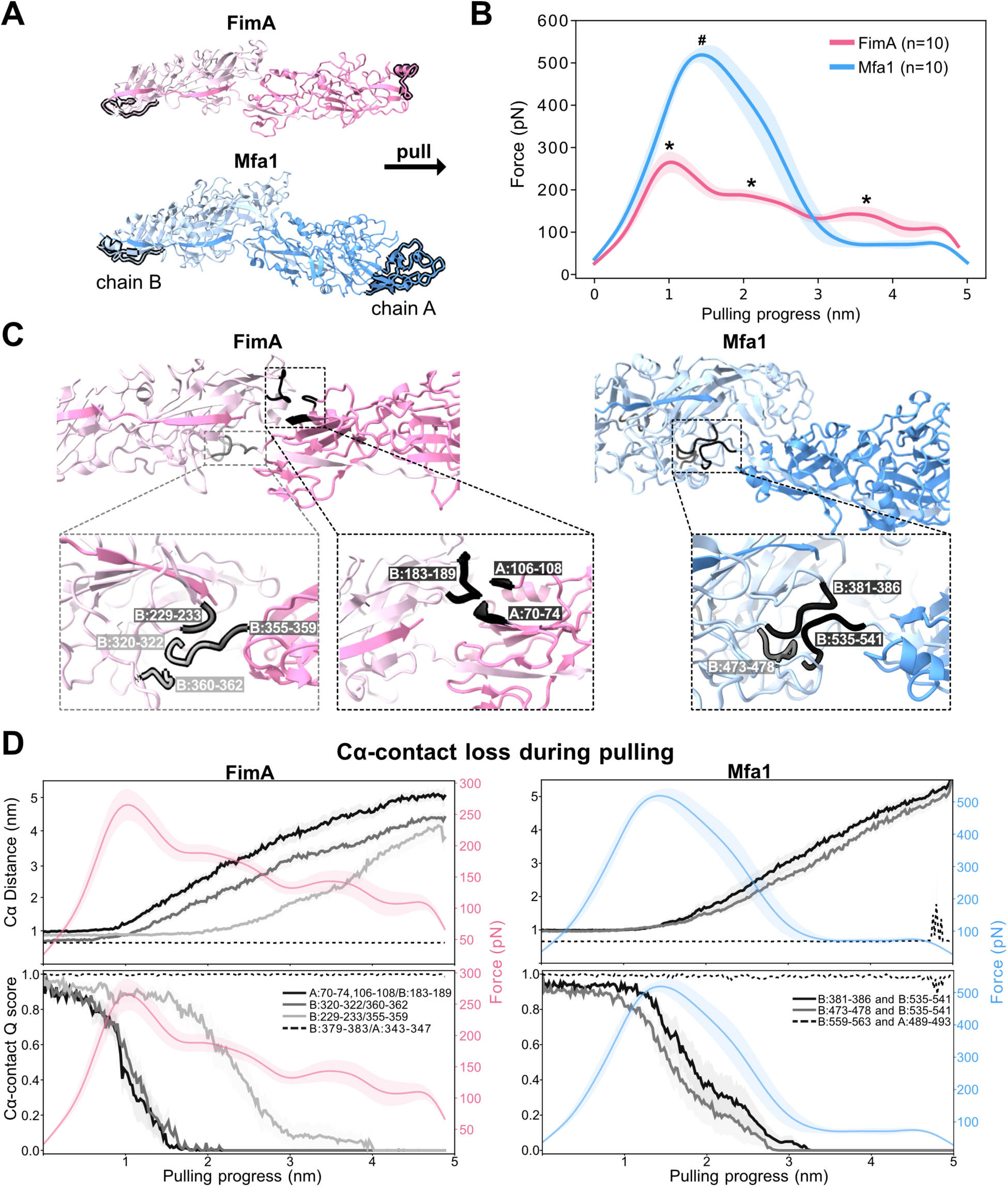
SMD simulations reveal distinct axial tensile responses in Fim and Mfa stalk fragments while donor-strand contacts remain engaged. **(A)** SMD simulations setup for Fim and Mfa stalk fragments composed of adjacent FimA or Mfa1 subunits, respectively. One end of each fimbriae fragment was anchored, and pulling force was applied to the opposite end along the longitudinal axis of the filament. Anchored and pulled groups are shown as black silhouettes, and the black arrow indicates the pulling direction. (**B**) Average force-extension profiles from ten SMD simulations of FimA and Mfa1 dimers at a pulling speed of 0.10 nm/ns. Lines represent the mean from replicates and shaded regions indicate standard error of the mean (SEM) across replicates. FimA showed a more distributed force-extension response across three distinct transition events (indicated by ‘**\***’), whereas Mfa1 exhibited a more concentrated response dominated by a single high-force transition (indicated by ‘**#**’). (**C**) Initial SMD structures highlighting selected Cα-contact regions that reproducibly lost proximity across all simulation replicates during pulling. In FimA, these regions include one inter-subunit contact involving residues 70-74 and 106-108 in subunit A and residues 183-189 in subunit B, as well as two intramolecular contacts within subunit B involving residues 320-322/360-362 and 229-233/355-359. In Mfa1, the monitored regions are intramolecular contacts within subunit B involving residues 381-386/535-541 and 473-478/535-541. Enlarged views highlight the corresponding contact regions in each stalk fragment. (**D**) Data are from ten independent simulations performed at a pulling speed of 0.10 nm/ns. Top panels show changes in Cα-Cα distances between selected residue pairs, with the corresponding force traces overlaid on the right y-axis. For each contact region, the plotted distance is the median of the eligible intergroup Cα-Cα distances; intrachain residue pairs separated by 20 or fewer positions in sequence were excluded. Dashed black traces show representative donor-strand-associated Cα-Cα distances (FimA, residues 379-383 in chain B and 343-347 in chain A; Mfa1, residues 559-563 in chain B and 489-493 in chain A), which remain largely unchanged during pulling, indicating that the exchanged donor strand remains engaged throughout the simulations. Bottom panels show normalized contact retention for the same regions, reported as the contact Q score, defined as the fraction of selected Cα contacts retained within 0.9 nm relative to the native Cα contacts present in the starting structure. Lines show the mean across simulations, and shaded bands indicate the SEM.

FimA and Mfa1 displayed distinct force-extension profiles. At 0.10 nm ns⁻¹, the mean profile across ten replicate trajectories showed multiple lower-force peaks for FimA, whereas the Mfa1 response was dominated by a higher-force peak (**Fig. 4B**). Analysis of the initial force peak across the five tested velocities showed that Mfa1 consistently required a higher force than FimA to traverse this transition at each matched pulling speed (**Fig. S4**). Thus, the two stalk fragments differed both in the forces associated with their initial transitions and in the distribution of subsequent force events along the extension trajectory.

To relate these force profiles to structural events, we first identified residue pairs that reproducibly lost native contacts near the principal force peaks in simulations performed at 0.10 nm ns⁻¹. We then tracked the median Cα-Cα distances and retained native Cα-contact fractions for the selected regions across replicate trajectories and pulling velocities (**Fig. 4C, D; Fig. S5**). This analysis distinguished changes in local contact networks from retention of the donor-strand-mediated connection between adjacent pilins.

In both systems, the exchanged C-terminal donor strands, spanning residues 363-383 in FimA and 543-563 in Mfa1, remained engaged in their acceptor grooves over the simulated extension range.

Representative donor-strand-associated contact pairs retained proximity while neighboring contacts separated (**Fig. 4D; Fig. S5; Movies S3 and S4**). The principal contact-loss events instead involved regions preceding the donor strands and adjacent inter-subunit or intra-subunit contacts. Axial deformation therefore occurred without extraction of the exchanged strand.

The distribution and sequence of these rearrangements differed between FimA and Mfa1. In FimA, the first major force peak coincided with disruption of both inter-subunit and intra-subunit contacts. The inter-subunit network involved residues 70-74 and 106-108 from one stalk pilin and residues 183-189 from the neighboring subunit. An intra-subunit contact between residues 320-322 and 360-362 was also disrupted during the early response (**Fig. 4C, D; Fig. S5**). At a later stage of extension, contact was lost between residues 229-233 and 355-359 within a pilin subunit. FimA therefore accommodated extension through sequential rearrangements involving both non-donor-strand contacts between adjacent pilins and contacts within the pilin fold.

In Mfa1, the principal monitored contact-loss events occurred within an intra-subunit network comprising residues 381-386, 473-478, and 535-541 (**Fig. 4C, D; Fig. S5**). The two monitored contact pairs, 381-386/535-541 and 473-478/535-541, lost proximity over overlapping extension ranges associated with the dominant force transition. Both pairs shared the 535-541 segment immediately preceding the exchanged donor strand. Thus, the mapped Mfa1 response centered on rearrangement of this donor-strand-adjacent intra-subunit network, in contrast to the combination of inter-subunit and intra-subunit transitions observed in FimA.

To determine whether the regions participating in these force-induced rearrangements are conserved across pilin genotypes, we mapped the SMD-defined contact regions onto the structure-guided sequence alignments described above (**Figs. S6, S7**). Notably, several of the contact regions that reorganized during stretching contained highly conserved residues despite the broader sequence diversity within each pilin family. In FimA, the early-disrupted 70-74/106-108/183-189 inter-subunit network and the later-disrupted 229-233/355-359 intra-subunit network contained highly conserved residues across the five different genotypes (**Fig. S6)**. By contrast, the early-disrupted 320-322/360-362 network contained conserved positions interspersed with genotype-associated substitutions (**Fig. S6**). Sequence conservation therefore extended to multiple FimA regions involved in both early and later stages of axial deformation, although it was not uniform across all responsive contact regions.

A corresponding relationship was observed in Mfa1. The 381-386 and 535-541 segments associated with the dominant force transition were highly conserved across the 70A, 70B, and 53 representatives examined. The additional interacting segment, residues 473-478, was also broadly conserved but contained limited group-associated substitutions (**Fig. S7**). Thus, in both pilin families, several of the regions that underwent force-induced contact rearrangements were sequence-conserved rather than confined to genotype-variable segments. These correspondences place conserved sequence elements within the deformation pathways identified for the ATCC 33277-derived stalks.

Together, the simulations distinguish retention of donor-strand-mediated inter-subunit attachment from rearrangement of the surrounding contact networks. FimA accommodates axial extension through sequential inter-subunit and intra-subunit transitions, whereas Mfa1 exhibits a dominant higher-force transition associated primarily with intra-subunit contact loss. Importantly, several regions participating in these distinct transitions are conserved across genotypes within their respective pilin families. The combined SMD and sequence analyses therefore identify conserved local elements that undergo rearrangement as the stalks extend, linking sequence conservation to the molecular deformation pathways surrounding the retained donor-strand connection.

### Native FimA and Mfa1 stalks differ in apparent local stiffness

AFM force mapping has been used to relate bacterial surface nanotopography to spatial variations in the apparent stiffness of cell envelopes and associated extracellular material^36,37^. To complement the axial pulling simulations, we used AFM force mapping to compare the resistance of purified nFim and nMfa filaments to local tip-induced indentation. Representative AFM height images demonstrated the same broad fimbriae length differences observed in the cryo-EM micrographs (**Fig. 5A**). The mean apparent Young’s modulus was 121.2 MPa for nFimA and 52.1 MPa for nMfa1, corresponding to a ∼2.3-fold difference (**Fig. 5B-D**). These measurements indicate greater local resistance to indentation in the FimA stalk under the conditions examined.

**Figure 5.**
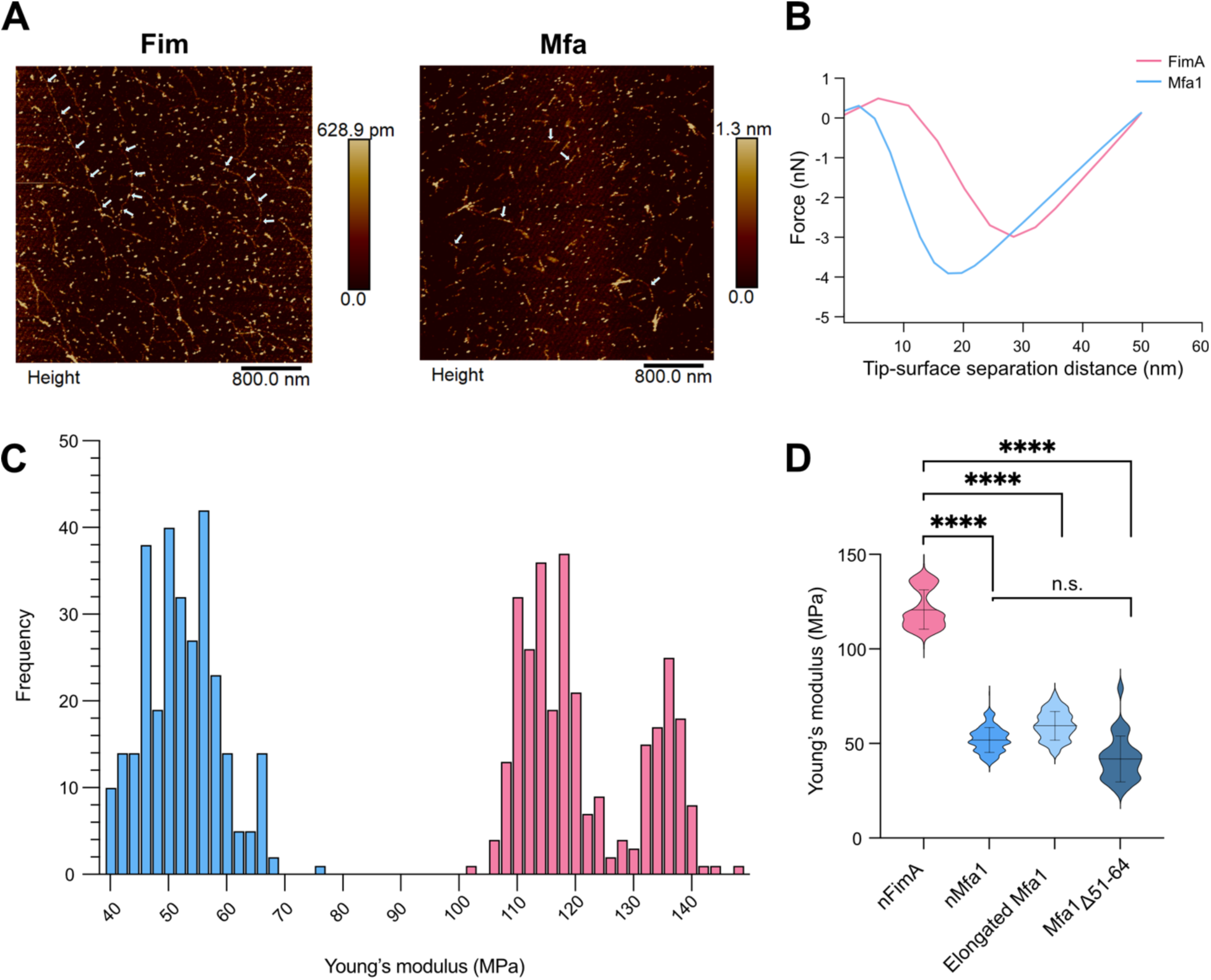
Native Fim fimbriae are locally stiffer than Mfa fimbriae under AFM indentation. **(A)** Representative AFM height images of purified native Fim and Mfa fimbriae. Fim fimbriae appeared as long, extended filaments, whereas Mfa fimbriae were shorter under the same imaging conditions. White arrows indicate representative filaments. Multiple arrows trace the path of selected Fim filaments, whereas individual arrows mark representative shorter Mfa filaments. Scan area, 4 µm × 4 µm. **(B)** Representative force-distance curves collected from native FimA and Mfa1 fimbriae. Measurements were collected from central regions of individual filaments to minimize contributions from terminal accessory pilin components. **(C)** Distribution of Young’s modulus values measured for native FimA and Mfa1 fimbriae. Native FimA (pink) displayed a higher apparent Young’s modulus than native Mfa1 (blue), indicating greater local stiffness under AFM indentation. **(D)** Summary of Young’s modulus values for native FimA, native Mfa1, elongated Mfa1, and Mfa1_Δ51-64_ fimbriae. Elongated Mfa1 and Mfa1_Δ51-64_ remained more compliant than FimA and were not significantly different from native Mfa1 under the conditions tested. One-way ANOVA was used to calculate the statistics: **** P < 0.0001; n.s., not significant.

To assess whether filament length alone could account for the difference in apparent local stiffness, we analyzed elongated Mfa fimbriae purified from a complemented Δmfa1 background in which the downstream insertion disrupts expression of *mfa2*-*mfa5*, including the Mfa2 anchoring and length-regulatory protein. This preparation provided an elongated Mfa counterpart to the unusually long Fim filaments produced by ATCC 33277, whose elongation is associated with a naturally occurring mutation in *fimB*, encoding the corresponding FimB anchoring and length-regulatory protein^19^. The elongated Mfa filaments exceeded 5 µm in length, reaching or surpassing the lengths observed for nFim (**Fig. S8**). Their mean apparent Young’s modulus was 59.6 MPa, higher than the shorter nMfa1 preparation but significantly lower than nFimA (**Fig. 5D**). The persistence of lower apparent stiffness in elongated Mfa filaments therefore argues against filament length alone accounting for the FimA-Mfa1 difference.

Together, AFM and SMD distinguish local indentation stiffness from the forces associated with axial structural rearrangement. FimA exhibits greater apparent local stiffness under indentation but undergoes multiple lower-force transitions during simulated axial extension. Mfa1, by contrast, is more compliant under indentation yet exhibits a dominant higher-force transition during simulated axial pulling. Thus, the relative mechanical responses of the two stalks differ between the assays, rather than defining a uniform ranking of mechanical strength.

### Latch deletion reduces heat-resistant Mfa1 polymer accumulation without abolishing stalk assembly

To test the contribution of the Mfa1 N-terminal latch to polymer formation, stalk architecture, and mechanics, we generated an Mfa1 variant lacking residues 51-64 (Mfa1_Δ51-64_) and expressed it under the native *mfa* operon promoter in a *P. gingivalis* ATCC 33277 Δ*fimA*Δ*mfa1* background.

Whole-cell immunoblotting showed the expected high-molecular-weight ladder following wild-type *mfa1* complementation after heating at 80 °C under nonreducing conditions^10,24,25^ (**Fig. 6A**). In contrast, Mfa1_Δ51-64_ exhibited a marked reduction in heat-resistant oligomers and additional slower-migrating bands, potentially representing immature precursor forms that are incompletely processed (**Fig. 6, black arrows**). Mature-sized species were also detected, including monomers after heating at 95 °C under reducing conditions, and assembled mutant filaments could be purified (**Fig. 6A**).

**Figure 6.**
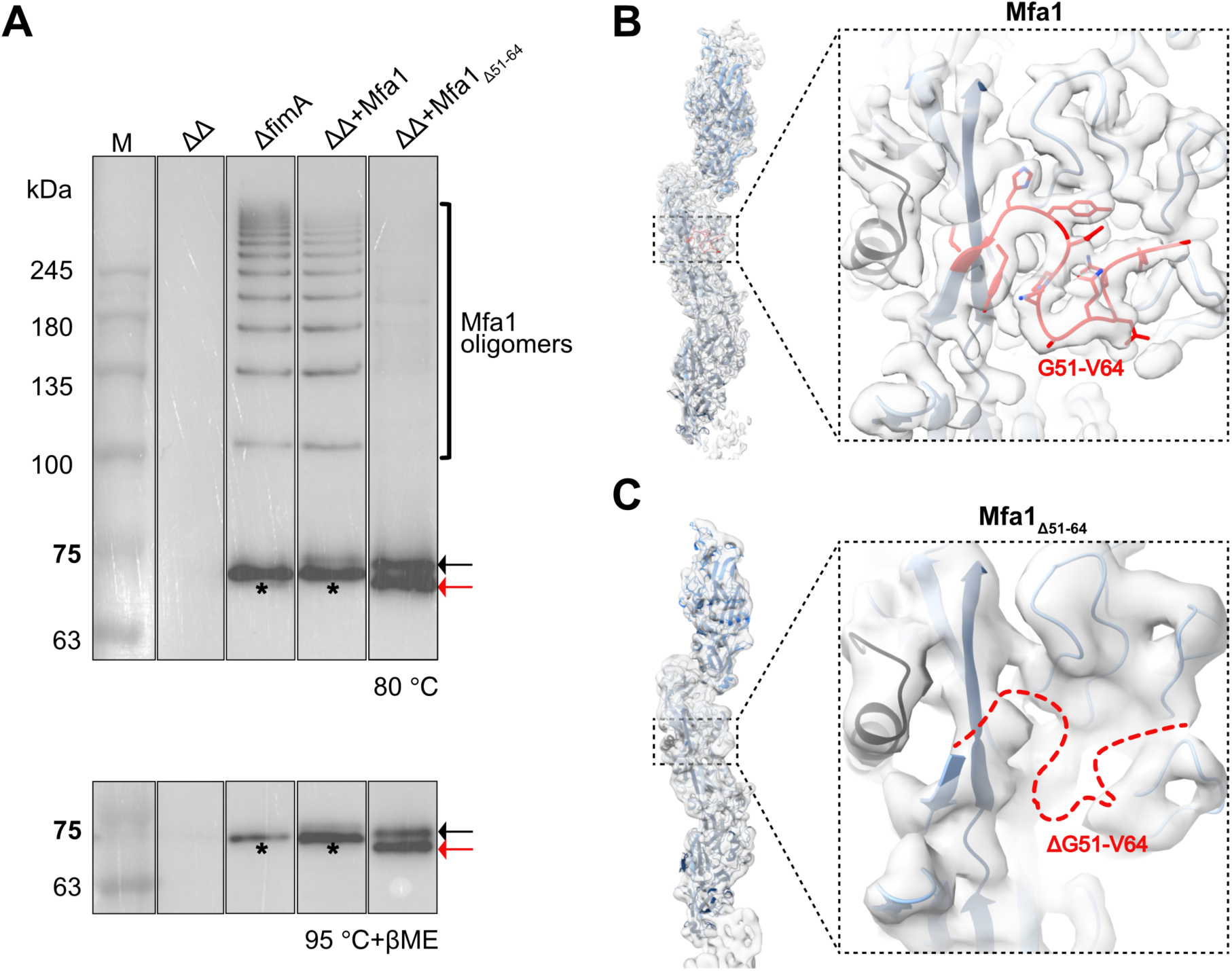
The Mfa1 N-terminal latch promotes heat-resistant Mfa polymer accumulation but is dispensable for the mature stalk architecture. (**A**) Anti-Mfa1 immunoblots of sonicated whole-cell samples after heat treatment at 80 °C under nonreducing conditions and at 95 °C with 10% β-mercaptoethanol (βME). High-molecular-weight ladder species correspond to Mfa1 oligomers, and the lower bands indicated by black asterisks correspond to monomeric Mfa1. Wild-type *mfa1* and *mfa1_Δ51-64_* were expressed from pT-COW under the native *mfa1* promoter in a *P. gingivalis ΔfimA::erm^R^Δmfa1::cm^R^* double mutant background (*ΔΔ)*. Lanes are” M, protein molecular weight marker; *ΔfimA*, single-mutant control expressing endogenous *mfa1*; *ΔΔ+Mfa1*, *ΔΔ* complemented with pT-COW-*np*-*mfa1*; *ΔΔ+*Mfa1_Δ51-64_, *ΔΔ* complemented with pT-COW-*np*-*mfa1_Δ51-64_*. Black arrows indicate slower-migrating Mfa1_Δ51-64_ species consistent with incompletely proteolytically processed or immature forms; red arrows indicate mature Mfa1_Δ51-64_. The presence of mature Mfa1_Δ51-64_ and recoverable assembled filaments indicate that deletion of residues 51-64 does not abolish polymerization, but the reduced high-molecular-weight ladder suggests decreased accumulation or heat resistance of Mfa1 polymers. **(B)** Cryo-EM map and fitted atomic model of the wild-type native Mfa1 stalk, shown across three successive Mfa1 stalk pilin subunits. The zoomed-in view highlights the Mfa1 N-terminal latch region spanning residues 51-64, which folds over the exchanged C-terminal donor strand. The latch region is outlined with a dashed red line, and the exchanged donor strand is shown in blue. **(C)** Cryo-EM map and fitted model of the Mfa1_Δ51-64_ stalk, reconstructed at 3.78 Å resolution and shown across three successive Mfa1_Δ51-64_ stalk fimbriae subunits. The zoomed-in view shows loss of density for the deleted latch region, in red dashed line, while the surrounding donor-strand-exchanged stalk architecture is retained. The overall similarity between the wild-type and Mfa1_Δ51-64_ stalks indicates that the N-terminal latch is not required to maintain the mature, polymerized fold once the stalk is assembled.

Cryo-EM analysis of the recovered mutant filaments yielded a reconstruction at 3.78 Å resolution. Docking of the native Mfa1 stalk model showed preservation of the overall pilin fold and donor-strand insertion, with no density at the position occupied by the deleted latch (**Fig. 6B, C**). The mutant nevertheless exhibited modest differences in helical packing, with an axial rise of 73.1 Å, a twist of 97.7°, and a pitch of 269.3 Å, compared with 71.7 Å, 92.4°, and 279.1 Å, respectively, for nMfa1 (**Figs. S2 and S9**). These values correspond to 3.7 subunits per helical turn in Mfa1_Δ51-64_, compared with 3.9 in nMfa1.

Surface-area analysis by PISA showed that latch deletion reduced the buried donor-strand surface by 251.7 Å², corresponding to a 9.3% decrease relative to native Mfa1. The buried fraction decreased from 85.8% to 77.5%, demonstrating increased donor-strand exposure in the recovered mutant stalks.

To assess the mechanical consequences of these structural changes, we examined the mutant filaments by AFM and SMD. AFM indentation yielded a mean apparent Young’s modulus of 42.1 MPa for Mfa1_Δ51-64_, compared with 52.1 MPa for nMfa (**Fig. 5D; Fig. S8C**). In matched SMD simulations, initial force peaks were comparable between the mutant and native Mfa1 across the five pulling velocities examined, with no statistically significant difference detected at any tested velocity (**Fig. S4**).

Together, the reduced oligomer signal and additional slower-migrating species support a model in which latch deletion impairs productive Mfa1 maturation and polymer accumulation. However, the recovered filaments show that reduced donor-strand shielding and altered helical packing remain compatible with stalk assembly and retention of high initial force peaks during simulated axial extension.

## Discussion

High-resolution structures of FimA and Mfa1 stalks assembled by and purified directly from *P. gingivalis* establish the molecular architectures of both principal type V fimbriae polymers after maturation in their native producer. Unlike assemblies generated through heterologous expression and *in vitro* polymerization, these filaments underwent processing and assembly in the pathogen before isolation. Their close correspondence with previously reported recombinant assemblies establishes that the polymerized pilin conformations defined in those systems are also present in pathogen-assembled stalks^24,25^. The native structures provide the experimental foundation for comparing the two systems and connecting their molecular organization to distinct mechanical responses. Together, these analyses show that a shared donor-strand exchange mechanism can support substantial variation in the organization and deformation of bacterial adhesive polymers.

The simulations distinguish maintenance of inter-subunit attachment from preservation of the resting stalk conformation. Exchanged donor strands remained engaged while neighboring inter-subunit and intra-subunit contacts rearranged, identifying different routes through which FimA and Mfa1 accommodate axial extension. This separation has parallels in other bacterial pili. In *Escherichia coli* type 1 fimbriae, reversible helical uncoiling permits extension while modulating the load transmitted to the terminal adhesin-receptor bond^28^. In *Acinetobacter baumannii* Csu pili, additional inter-subunit “clinch” contacts organize the zigzag architecture and constrain subunit movement during extension^38^. These examples illustrate how contacts organizing a filament can rearrange without immediately disrupting polymer connectivity. The type V simulations identify local changes within and between pilins rather than establishing reversible whole-filament uncoiling.

Several regions participating in these force-induced rearrangements contain residues conserved across genotypes within their respective pilin families (**Figs. S6, S7**). Their conservation is consistent with an important contribution to the interactions that resist and accommodate tensile loading. Preserving these contact regions may help maintain characteristic aspects of stalk mechanics despite broader sequence diversity, suggesting that mechanical requirements could contribute to their conservation alongside constraints imposed by folding and assembly. The correspondence between conservation and simulated contact rearrangement therefore supports a mechanical role for these local elements.

The latch deletion separates the possible contribution of the retained Mfa1 N-terminal region to maturation from its role in donor-strand shielding. Reduced heat/SDS-resistant oligomer abundance and additional slower-migrating species support a model in which deletion impairs productive processing, limiting the availability of assembly-competent subunits (**Fig. 6A**). Nevertheless, recovered filaments retained the donor-strand-exchanged architecture and comparable initial SMD force peaks despite reduced donor-strand burial. Thus, reduced polymer accumulation can coexist with preservation of the principal tensile signature in filaments that successfully assemble. This finding also argues against N-terminal shielding alone explaining the higher initial force response of Mfa1.

Assembly-associated remodeling provides a more specific connection to adhesive function. The repositioned Mfa1 α5 region lies within the segment implicated in recognition of the streptococcal adhesin SspB. Substitution of A357 within α5 with proline reduced adherence to *Streptococcus gordonii* while retaining surface expression and polymerization behavior similar to wild type^39^. Together with the structural comparison, this phenotype supports a contribution of local conformation to adhesive-surface presentation, without identifying A357 as a direct receptor-contacting residue. Predicted variation around α5 and the upper loops of assembly-competent Mfa1-53 further indicates that polymer formation accommodates sequence variation in regions potentially involved in adhesive presentation.

Since FimA and Mfa1 themselves participate in adhesion, stalk geometry may influence the spacing and orientation of interaction sites, while deformation may affect their engagement during displacement between attached cells or surfaces^9,40^. A direct contribution of stalk architecture to community organization is illustrated by Csu pili, whose rods form lateral junctions and antiparallel stacks connecting bacteria within biofilms, distinct from tip-mediated surface attachment^41^. The relevant parallel is that the repeating stalk can organize adhesive contacts rather than simply support a terminal adhesin. Stalk mechanics may therefore complement receptor specificity by influencing the physical behavior of an established connection.

We propose that the distinct FimA and Mfa1 responses could contribute to their partly overlapping adhesive roles. For FimA, greater local indentation resistance combined with sequential lower-force axial transitions is consistent with resistance to local deformation alongside routes for progressive extension. Such properties could help maintain adhesive-surface organization during engagement with diverse substrates while accommodating displacement between binding partners. Mfa1 combines greater local compliance under indentation with a higher force requirement for its principal axial transition. This combination could favor local accommodation at cell–cell interfaces while resisting axial rearrangement, potentially supporting close interbacterial contacts.

These interpretations offer a possible physical contribution to the broader adhesive engagement associated with Fim and the prominent coadhesive role of Mfa, without assigning either system an exclusive function.

The elongated Mfa1 filament retained a lower apparent indentation modulus than FimA, arguing against filament length alone explaining the measured difference. Nevertheless, anchoring and length regulation influence filament reach, and the intrinsic *fimB* mutation in ATCC 33277 limits generalization of its elongated Fim phenotype^19^. The biological model also remains bounded by the assays: AFM measures surface-supported filaments under the reported preparation conditions, whereas SMD resolves transitions in short fragments at imposed pulling rates. These complementary comparisons do not isolate loading direction as the only variable or directly measure adhesive performance in biofilms.

Together, the native-source structures and mechanical analyses identify distinct deformation pathways surrounding a shared donor-strand linkage. The findings support a model in which conserved local contacts contribute to tensile responses, while donor-strand shielding and productive polymer formation have distinguishable roles. This molecular organization provides a basis for understanding how stalk mechanics may complement ligand recognition in sustaining the different adhesive interactions supported by Fim and Mfa.

## Materials and Methods

### *P. gingivalis* fimbriae isolation and purification

#### Bacterial strains and culture conditions

*P. gingivalis* ATCC 33277 wild-type and variant strains were grown on Trypticase Soy (TSB) Agar plates (Becton, Dickinson and Company, Franklin Lakes, NJ, USA) supplemented with 5 µg/mL hemin, 1 µg/mL menadione and 5% defibrinated sheep blood (BAPHK) (HemoStat Laboratories, USA) at 37 °C for 3-5 days under anaerobic conditions (5% hydrogen, 10% carbon dioxide and 85% nitrogen). Overnight liquid cultures were prepared by inoculating pigmented colonies into TSBHK medium and incubating anaerobically at 37 °C without agitation. Appropriate antibiotics were supplemented as required.

#### Purification of major Fim fimbriae and elongated minor Mfa fimbriae

Native Fim fimbriae were purified from *P. gingivalis* ATCC 33277 Δ*mfa1::erm^R^* strain cells using an adapted version of a previously described protocol for T-pilus purification. Briefly, bacterial cells were harvested by centrifugation at 5,000 x g for 15 min, washed with phosphate-buffered saline (PBS), and resuspended in 20 mM Tris-HCl, pH 8.0, 150 mM NaCl, and 10 mM MgCl₂. Fim fimbriae were mechanically sheared from the cell surface by magnetic stirring for 30 min at room temperature. Cell debris was removed by centrifugation, and crude fimbriae were pelleted by ultracentrifugation at 150,000 × g for 4 hours at 4 °C. The pellet was resuspended in 20 mM Tris-HCl pH 7.5, 150 mM NaCl, and 0.5% sodium deoxycholate. Solubilized fimbriae were further purified by sucrose velocity sedimentation on a 5-50% sucrose gradient. Fractions containing Fim fimbriae were pooled, and sucrose was removed by repeated buffer exchange in a 100-kDa molecular-weight-cutoff (MWCO) centrifugal filter device (Amicon^®^). The elongated Mfa fimbriae were similarly purified from *P. gingivalis* ATCC 33277 *ΔfimA::cm^R^Δmfa1::erm^R^ +* pT-COW-*np-mfa1* strain.

#### Purification of minor Mfa fimbriae

Native Mfa fimbriae were purified from *P. gingivalis* ATCC 33277 Δ*fimA::erm^R^*strain cells as previously described^42^. Briefly, bacterial cells from liquid culture were harvested by centrifugation at 5,000 x g for 15 min, washed with PBS, and resuspended in 10 mM HEPES pH 7.4 supplemented with EDTA-free protease inhibitor cocktail (Millipore). Cells were lysed by using a French press at 100 MPa and 4 °C and cell debris was removed by ultracentrifugation at 100,000 × g for 1 hour at 4 °C. Ammonium sulfate was added to the soluble fraction to 50% saturation, and the precipitated proteins were pelleted by centrifugation at 15,000 x g for 30 min. The protein pellet was resuspended in 20 mM Tris-HCl pH 8.0 and dialyzed against the same buffer overnight. Native Mfa fimbriae were further purified by ion-exchange chromatography using 12-mL DEAE Sepharose™ Fast Flow resin (Cytiva), followed by size exclusion chromatography on a Superose 6 10/300 GL column (Cytiva) pre-equilibrated in 20 mM Tris pH 8, and 150 mM NaCl.

#### Negative stain and transmission electron microscopy

Purified fimbriae samples were diluted to 0.05 mg/mL, and 5 μL of each sample was applied to carbon-coated electron microscopy copper grids that had been glow-discharged for 30 s at 10 mA. After 5 min incubation, excess sample was blotted away. The grids were washed with ddH_2_O and subsequently stained with 1% (w/v) uranyl acetate for 1 min. Excess stain was blotted, and grids were air-dried. Negatively stained fimbriae were imaged using a FEI Tecnai G2 Spirit BioTwin transmission electron microscope operated at an accelerating voltage of 120 kV.

### Single-particle analysis on purified fimbriae

#### Cryo-EM sample preparation and data collection

Purified fimbriae were applied to glow-discharged (Pelco easiGlow Glow Discharge Cleaning System, Negative) holey carbon grid (C-flat CF-2/1-3Cu-T; 10 mA for 10 s), blotted and vitrified using a Vitrobot Mark IV (Thermo Fisher Scientific) at 4°C and 100% humidity. For nFim sample, 3.5 µL of sample at 0.5 mg/mL was applied to the grid and blotted for 2 s at blot force 3 prior to vitrification. For nMfa and nMfa1_Δ51-64_ sample, 3.5 µL of sample at 0.15 mg/mL was applied and blotted for 3 s at blot force 1 prior to vitrification. For nFim and nMfa, Cryo-EM movies were collected using the SerialEM software (v4.1.0β)^43^ on a Titan Krios transmission electron microscope (Thermo Fisher Scientific) equipped with a Gatan K3 direct electron detector and a Quantum LS energy filter at the Facility for Electron Microscopy Research (FEMR), McGill University. For nMfa1_Δ51-64_, Cryo-EM movies were collected using the E Pluribus Unum (EPU) software on a Thermo Scientific Glacios 2 cryo-transmission electron microscope operated at 200 kV and equipped with a Falcon 4i direct electron detector and a Selectris energy filter at FEMR McGill University. Data collection parameters are summarized in **Table S1**.

#### Cryo-EM data processing and model building

Cryo-EM data processing was performed in CryoSPARC v4^44^. Movies were aligned and corrected for beam-induced motion using patch motion correction. Contrast transfer function (CTF) parameters were estimated and corrected using patch CTF estimation. Particles were picked using a Filament tracer without templates with a maximum diameter of 45 Å and 57 Å for FimA and Mfa1, respectively. Particles were extracted using box sizes of 480 and 640 pixels for FimA and Mfa1, respectively. Extracted particles were subjected to iterative rounds of 2D classification, and selected classes were used for helical reconstructions.

For FimA, selected particles were subjected to helical reconstruction. Initial volumes were generated without imposed symmetry and used to estimate the initial helical parameters. Following helical refinement, reference-based motion correction was performed to correct per-particle beam-induced motion, followed by local refinement and local CTF refinement to obtain the final reconstruction. For Mfa1, helical reconstruction using the corresponding helical parameters yielded a map at 3.64 Å resolution. In parallel, selected Mfa particles were subjected to *ab initio* reconstruction followed by non-uniform and local refinement, which improved the reconstruction to 2.9 Å resolution and was therefore used as the final Mfa map for structural analysis. Final map resolutions were estimated using the gold-standard Fourier shell correlation (FSC) criterion at the 0.143 threshold. Processing workflows for both datasets are shown in **Figure S1**.

#### Model building and structural validation

Initial models were generated in Coot using the previously reported FimA and Mfa1 structures, PDB entries 6JZK^25^ and 5NF2^10^, respectively. Template models were fitted into the cryo-EM maps in ChimeraX^45^ and regions that were unresolved in the template structures were manually built into the density in Coot^46^. The models were refined iteratively through manual rebuilding in Coot^46^ and real-space refinement in PHENIX with non-crystallographic symmetry restraints applied^47^. Model quality was assessed using MolProbity^48^. Structural deposition and refinement statistics are summarized in **Table S1**.

### Thermal stability assay of P. gingivalis *fimbriae* in whole cells

#### Generation of deletion strains in *P. gingivalis*

The two *P. gingivalis* ATCC33277 double mutant strains were generated as previously described^49^. The *P. gingivalis* ATCC 33277 *ΔfimA::cm^R^Δmfa1::erm^R^*strain was generated in the *Δmfa1::erm^R^* background by replacing *fimA* with a chloramphenicol-resistance cassette (*cm^R^*). Conversely, the ATCC 33277 *ΔfimA::erm^R^Δmfa1::cm^R^* strain was generated in the *ΔfimA::erm^R^* background by replacing *mfa1* with *cm^R^*. Briefly, ∼1-kb regions upstream and downstream of the *fimA or mfa1* gene were amplified from *P. gingivalis* ATCC 33277 genomic DNA and a *cm^R^* cassette that was cloned from the pACYC-Duet vector. The three amplicons were then assembled to generate a linear deletion construct in which the *cm^R^* flanked by the *fimA or mfa1* upstream and downstream homology regions. The resulting linear DNA fragment was used to transform *P. gingivalis* ATCC33277 *Δmfa1::erm^R^*or *ΔfimA::erm^R^* cells, and transformants were selected on BAPHK supplemented with chloramphenicol (10 μg/mL). Primers and plasmids used in this study are listed in **Tables S2** and **S3**, respectively.

#### Complementation in *P. gingivalis*

The wildtype *mfa1* gene together with its native promoter (np) region were amplified from *P. gingivalis* ATCC33277 and cloned into the pT-COW *E. coli-P. gingivalis* shuttle plasmid to generate pT-COW-*np-mfa1*. The Mfa1 N-terminal deletion variant, lacking residues G51-V64, was generated by PCR-based site-directed mutagenesis. Plasmids were introduced into *P. gingivalis* by conjugation from *E. coli* S17-1^50^ via solid-surface mating, as previously described^51^. Briefly, donor *E. coli* S17-1 and recipient *P. gingivalis ΔfimA*Δ*mfa1* strains were grown overnight in TSBHK under aerobic and anaerobic conditions, respectively, to stationary phase. Equal volumes of donor and recipient cultures were combined and harvested by centrifugation. The pellets were resuspended in 100 µL TSBHK and spotted on agar plates. Plates were incubated aerobically at 37 °C for 3 hours to facilitate mating, then transferred to anaerobic conditions for further incubation and selection on BAPHK plates supplemented with tetracycline (1 μg/mL).

### Thermal stability assay

Overnight *P. gingivalis* cultures were harvested and normalized to an OD_600_ of 1.0 in PBS. Equal volumes of sonicated normalized cell suspension were mixed with Sodium Dodecyl Sulfate (10% SDS, 50% Glycerol, 500 mM Tris-HCl and 0.5% bromophenol blue dye) sample buffer and subjected to one of two heat treatments. For detection of heat-resistant Mfa1 oligomers, samples were heated at 80 °C for 5 min, under non-reducing conditions. For complete dissociation of Mfa1 fimbriae into monomers, samples were heated at 95 °C for 5 min in SDS sample buffer supplemented with 10% β-mercaptoethanol (βME). Samples were resolved by SDS-PAGE and analyzed by immunoblotting using an anti-Mfa1 antibody.

### Anti-Mfa1 antibody production

The gene encoding the mature *P. gingivalis* ATCC 33277 Mfa1 (residues 51-563) was cloned into a modified pET28a (+) plasmid incorporating an N-terminal His_10_-tag by Gibson assembly. All primers and plasmids are listed in **Tables S2 and S3**, respectively.

*E. coli* BL21 (DE3) cells harbouring pET28a-*mfa1* were grown in LB medium supplemented with kanamycin to an OD_600_∼0.5 at 37 °C. Recombinant protein expression was induced with Isopropyl β-D-1-thiogalactopyranoside (IPTG) and cultures were maintained at 20°C for an additional 16 h. Cells were harvested by centrifugation at 6,000 × g for 15 min at 4 °C. The cell pellet was resuspended in 20 mM Tris pH 8, 300 mM NaCl, 10 mM imidazole supplemented with DNase I (10 μg/ml) and protease inhibitor cocktail (Calbiochem) and lysed by sonication at 4°C. Cell debris was removed by ultracentrifugation at 270,000 × *g* for 1 hour at 4°C. The supernatant was applied onto a gravity-flow Ni-NTA column (Bio-Rad Econo-Column chromatography column, Thermo Scientific HisPur Ni-NTA resin). The bound protein was washed with buffers 20 mM Tris pH 8, 300 mM NaCl, 40 mM imidazole and eluted with buffer 20 mM Tris, 300 mM NaCl, 300 mM imidazole. Affinity tag was removed by bovine thrombin (Prolytix) digestion during an overnight dialysis against buffer 20 mM Tris at 4°C. Recombinant Mfa1 was further purified using size exclusion chromatography (HiLoad 16/600 Superose 6 column, Cytiva) in buffer 20 mM Tris pH 8, 200 mM NaCl, 2 mM β ME. The purified protein was concentrated to 1 mg/mL and flash-frozen in liquid nitrogen and was used as the immunogen for production of a custom rabbit polyclonal anti-Mfa1 antibody (Cederlane Laboratories).

### Atomic force microscopy (AFM)

The local stiffness of purified nMfa and nFim fimbriae was evaluated by AFM nano-indentation using an AFM Multimode-8 HR instrument (Bruker) operated with NanoScope 9.7 software. Imaging was performed in the PeakForce quantitative nanomechanical mapping (QNM^TM^) tapping mode capturing simultaneously data of channels “Height” and “DMT Modulus” to render images of surface topography and Young’s modulus, respectively. Samples were prepared by depositing a drop of 200 µL buffer (20 mM Tris pH 8, and 150 mM NaCl) on freshly cleaved muscovite mica grade V4 (Electron Microscopy Sciences) substrates forming a liquid dome over the mica surface, followed by injection of 2 µL of freshly prepared fimbriae at 0.75 mg/mL. Samples were incubated in air in ambient conditions (22 °C) for 5 min, followed by pipetting off the liquid from the mica surface, then washed immediately with ddH_2_O while retaining visibly thin water layer. Each mica substrate was then immediately mounted on the AFM for measurement. Imaging was performed under ambient air, with the acquisition time limited to a maximum of 5 min per sample to minimize dehydration.

The AFM probes were ScanAsyst-Air, consisting of a silicon tip on silicon nitride cantilever, nominal spring constant *k* = 0.4 N/m, tip radius at apex = 1 nm. Imaging was performed using the instrument’s ScanAsyst Auto Control feedback with PeakForce Amplitude of 150 nm, PeakForce Frequency of 2 kHz, and Scan Rate of 2 Hz. Before scanning, the probes were calibrated for deflection sensitivity with sapphire (very hard, E =400 GPa) reference caliber resulting in sensitivities in the range of 30-40 nm/V. The cantilever spring constant k was determined by thermal tuning and ranged from 0.38 to 0.45 N/m. The calibrated spring constant and nominal radius at apex were used as input probe parameters for Young’s modulus determination. A Poisson’s ratio of 0.5 was assumed for the fimbriae, consistent with values commonly used for proteinaceous materials.

Samples were scanned on surface areas of 2 µm x 2 µm, 4 µm x 4 µm to 10 µm x 10 µm at a resolution of 512 points per line along the x-axis and 512 lines along the y-axis. Force-distance curves were recorded at each image pixel using the instrument’s “Enable PeakForce Capture” function. The apparent Young’s modulus (E) of the fimbriae was obtained from the DMT Modulus channel generated automatically by the AFM software by fitting the force–distance curves to the Derjaguin-Muller-Toporov (DMT) model. Fitting was performed on the unloading (retraction) portion of the force curve over the range corresponding to 20-90% of the maximum force.

Acquired images were processed using NanoScope Analysis 2.0 software (Bruker). Height images were digitally flattened, and brightness, contrast, z-axis scale, and z-axis offset were adjusted for both height and modulus images. Measurements of the Young’s modulus (E) of fimbria were taken in the middle of nMfa fimbriae (90-120 nm length) and in locations spaced at least 200 nm apart of nFim (0.5-2 µm length). For each fimbriae sample, three independently prepared mica surfaces were examined. Approximately 100 manually selected modulus measurements were obtained from each mica surface, yielding approximately 300 measurements per sample. Sampling positions were distributed across multiple spatially separated filaments within each imaged population rather than concentrated on a single filament. Because more than one measurement could be obtained from the same filament, individual modulus values were treated as technical measurements rather than independent biological replicates.

### Steered molecular dynamics (SMD) simulations

SMD simulations were designed as a comparative assay to examine how matched FimA, Mfa1, and Mfa1_Δ51-64_ stalk fragments respond to axial tensile loading, rather than to estimate absolute physiological rupture forces. SMD simulations were performed in GROMACS (version 2024)^52^ using FimA and Mfa1 stalk fragments containing two donor-strand exchanged subunits designated chains A and B. An additional terminal donor strand was included to occupy the acceptor groove of the distal subunit and preserve the donor-strand-exchanged interface at that end of the model, designated chain C. The donor strand of the pulled terminal subunit (chain A) was removed to establish a defined pulling geometry and reduce the required simulation box size.

System topologies were generated with the CHARMM36-jul2022 force field^53^ and solvated using the TIP3P water model^54^. Each fragment was aligned along its principal axis, centered in a triclinic box, solvated, and neutralized with Na^+^ and Cl^-^ ions to a final concentration of 0.15 M. Following energy minimization, systems were equilibrated through alternating restrained NVT and NPT stages in which the temperature was progressively increased from 100 K to 303.15 K while positional restraint force constants were gradually reduced. Each temperature stage consisted of a 200-ps NVT step followed by a 500-ps NPT step, using restraint force constants of 1000, 500, 200, 100, and 50 kJ·mol⁻¹ nm⁻² at 100, 150, 200, 250, and 303.15 K, respectively. This was followed by a final 10 ns unrestrained NPT equilibration step at 303.15 K. Temperature was maintained using the V-rescale thermostat^55^, and pressure during NPT equilibration was maintained at 1 bar using the C-rescale barostat^56^.

For production SMD runs, the proximal end of each stalk fragment was position-restrained (FimA: chain B, res: 135-150; Mfa1: chain B, res: 205-230), and force was applied to the distal end (FimA: chain A, res: 285-300; Mfa1: chain A, res: 249-271 and 394-458) along the longitudinal axis of the filament using the GROMACS pull code.

Pulling simulations were performed in the NVT ensemble at 303.15 K with umbrella pulling, over a total pulling distance of 5 nm using five pulling velocities: 0.02, 0.05, 0.10, 0.20, and 0.50 nm·ns^-1^. A spring constant of 500 kJ·mol^-1^ nm^-2^ was used for all simulations. Pull-coordinate and pull-force outputs were recorded for downstream force-extension analysis. The source code, configuration templates, and analysis utilities are publicly available at https://github.com/Adwaith99/fimbriae-stretch.

Since SMD pulling velocities are substantially faster than most experimental force spectroscopy measurements, force values were interpreted comparatively rather than as absolute physiological rupture forces. Analyses therefore focused on relative differences among matched FimA, Mfa1, and Mfa1_Δ51-64_ models and on reproducible contact loss events across replicate simulations.

### Cα-based measurements and analysis

Candidate force-sensitive regions were first identified from SMD trajectories collected at 0.10 nm·ns^-1^, the midpoint of the tested pulling-speed range. Native contacts were defined as pairs of protein non-hydrogen atoms separated by no more than 0.45 nm in the starting structure. During pulling, that contact was considered retained when the same atom-atom distance remained no greater than 0.60 nm. Intra-subunit contacts between residues separated by 20 or fewer positions in sequence were excluded. Residue pairs containing at least three non-hydrogen atom contacts were examined for contact loss near the major force peaks. Regions showing recurrent contact loss across replicate trajectories were selected for subsequent Cα-based analysis. This contact screen was used only to identify regions for visualization and was not treated as a statistical test.

The selected FimA regions were an inter-subunit region comprising chain A residues 70-74 and 106-108 versus chain B residues 183-189; an early intra-subunit region comprising chain B residues 320-322 versus 360-362; and a later intra-subunit region comprising chain B residues 229-233 versus 355-359. The selected Mfa1 regions comprised chain B residues 381-386 versus 535-541 and chain B residues 473-478 versus 535-541. Regions outside the dominant force-sensitive residue pairs comprised chain B residues 379-383 versus chain A residues 343-347 for FimA and chain B residues 559-563 versus chain A residues 489-493 for Mfa1.

Trajectories were analyzed using MDAnalysis 2.0^57^. Protein coordinates were sampled every 0.10 ns. The SMD pull coordinates were obtained from the GROMACS pull_pullx.xvg output. This coordinate is calculated from the center-of-mass separation between anchor and pulled atom groups. The pulling-progress axis therefore represents the magnitude of displacement of the GROMACS pull coordinate from the initial structure.

Two Cα-based measurements were calculated for each selected region. First, the median Cα distance was defined as the median of the Cα-Cα distances among the evaluated residue pairs. Second, the retained native Cα-contact fraction, Q, was defined as the fraction of Cα residue pairs that identified as native contacts in the starting structure within 0.90 nm:

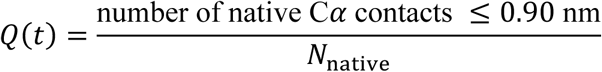

N_native_= total number of native Cα contact pairs in the starting structure;

Q=1 means all native contacts remain;

Q=0 means no native contacts remain.

For each speed, curves show the replicate means with standard error of the mean (SEM) shading, based on ten trajectories per system at each pulling speed. Force-extension traces were smoothed using discrete convolution with normalized one-dimensional Gaussian kernels implemented in NumPy (version 2.4.3)^58^, with half-widths at half-maximum (HWHM) of 5 ps along time and 0.20 nm along the extension.

### Intact protein liquid chromatography-mass spectrometry (LC-MS)

As previously described^59–61^, purified Mfa1 (residue 50-563) samples were diluted to 0.1 mg/mL in the presence of dithiothreitol (10 mM DTT final) and formic acid (0.1% (v/v) FA final) before 1 µg was injected on a Dionex Ultimate 3000 UHPLC system at 200 µL/min using a Waters BioResolve RP mAb Polyphenyl column (450Å, 2.7 µm, 2.1 x 100 mm). The resulting eluate (5 min wash with 4% (v/v) acetonitrile in 0.1% (v/v) formic acid, followed by 20 min gradient to 90% (v/v) acetonitrile in 0.1% (v/v) formic acid) was analyzed on an Impact II QTOF mass spectrometer (Bruker Daltonics) equipped with an Apollo II ion funnel electrospray ionization source and pre-calibrated (ESI-L Low Concentration Tuning Mix; Agilent Technologies #G1969-85000). Using the Bruker otofControl v4.0 / DataAnalysis v4.3 software, data were acquired in positive-ion profile mode using a capillary voltage of 4,500 V and dry nitrogen heated at 200 °C. For all intact samples (purified native Mfa fimbriae vs. commercial myoglobin and BSA standards), total ion chromatograms were used to determine where the proteins eluted, and spectra were summed over the entire elution peak. Multiply charged ion species were deconvoluted at a resolution of 5,000 using the maximum entropy method.

### Structure prediction, sequence alignment and phylogenetic analysis

The experimentally determined FimA-I and Mfa1-70A structures from *Porphyromonas gingivalis* ATCC 33277 were used as structural and residue-numbering references for comparisons across representative FimA genotypes I-V and the Mfa1-70A, Mfa1-70B, and Mfa1-53 groups. Structures of the remaining variants were predicted using AlphaFold3^62^, with a three-chain input configuration designed to favor intermolecular donor-strand complementation and the extended C-terminal donor-strand conformation observed in assembled fimbriae.

For each variant, the first input chain comprised the complete mature stalk pilin sequence beginning immediately after the assigned Rgp cleavage-site arginine. The second chain comprised the same mature sequence without its C-terminal donor strand, while the third consisted of the corresponding donor-strand sequence alone. This arrangement was designed to permit the C-terminal donor strand of the complete pilin to occupy the acceptor groove of the truncated neighboring subunit, while the isolated donor-strand segment complemented the groove of the complete pilin.

For each predicted assembly, only the chain containing the complete mature pilin sequence was retained for alignment. The FimA and Mfa1 datasets were aligned separately using FoldMason^33^, with the corresponding experimentally determined ATCC 33277 pilin included in each dataset. The resulting structure-guided amino acid alignments were visualized using ESPript 3.0^34,35^. Alignment numbering and secondary-structure annotations were referenced to the full-length ATCC 33277 FimA and Mfa1 precursor sequences, preserving the residue numbering used in the structural analyses.

Maximum-likelihood phylogenetic trees were inferred separately for the FimA and Mfa1 datasets using IQ-TREE 2^63^ and the corresponding structure-guided amino acid alignments.

## Acknowledgments

We thank Mark A. Hancock (McGill SPR-MS Facility) for assistance with the intact protein LC-MS analyses. The McGill SPR-MS Facility acknowledges infrastructure support from the Canada Foundation for Innovation (CFI) and the Centre de Recherche en Biologie Structurale (CRBS). We thank K. Basu and K. Sears at the Facility for Electron Microscopy Research at McGill University for assistance with microscope operation and cryo-EM data collection. FEMR is supported by the Canada Foundation for Innovation, the Government of Quebec, and McGill University. Molecular dynamics simulations were supported by resources from the Digital Research Alliance of Canada.

## Authors contributions

Z.W. and N.Z. initiated the project. Z.W. performed cryo-EM sample preparation, data processing, model building, biochemical experiments, structural analysis, steered molecular dynamics simulations, and visualization. A.B.U. and D.P.T contributed to steered molecular dynamics simulation design and analysis. V.D.N. and D.P.R. performed the AFM experiments and analysis. Z.W., A.B.U., V.D.N., D.P.R., D.P.T., K.H.B., and N.Z. contributed to investigation and formal analysis. K.H.B., D.P.R., and N.Z. supervised the work. N.Z. and K.H.B. acquired funding. Z.W. and N.Z. wrote the original draft. All authors reviewed and edited the manuscript.

## Data availability

Electron microscopy density maps have been deposited in the Electron Microscopy Databank (EMD-78109, EMD-78110, and EMD-78111). Atomic models have been deposited in the Protein Databank (37EB, 37EC, and 37ED).

## References

(1) Mei, F.; Xie, M.; Huang, X.; Long, Y.; Lu, X.; Wang, X.; Chen, L. Porphyromonas Gingivalis and Its Systemic Impact: Current Status. Pathog. Basel Switz. 2020, 9 (11). 10.3390/pathogens9110944

(2) Honda, K. Porphyromonas Gingivalis Sinks Teeth into the Oral Microbiota and Periodontal Disease. Cell Host Microbe 2011, 10 (5), 423–425. 10.1016/j.chom.2011.10.008

(3) Bostanci, N.; Belibasakis, G. N. Porphyromonas Gingivalis: An Invasive and Evasive Opportunistic Oral Pathogen. FEMS Microbiol. Lett. 2012, 333 (1), 1–9. 10.1111/j.1574-6968.2012.02579.x

(4) Holt, S. C.; Ebersole, J. L. Porphyromonas Gingivalis, Treponema Denticola, and Tannerella Forsythia: The “Red Complex”, a Prototype Polybacterial Pathogenic Consortium in Periodontitis. Periodontol. 2000 2005, 38, 72–122. 10.1111/j.1600-0757.2005.00113.x

(5) Bao, K.; Belibasakis, G. N.; Thurnheer, T.; Aduse-Opoku, J.; Curtis, M. A.; Bostanci, N. Role of Porphyromonas Gingivalis Gingipains in Multi-Species Biofilm Formation. BMC Microbiol. 2014, 14, 258. 10.1186/s12866-014-0258-7

(6) Lamont, R. J.; Kuboniwa, M. The Polymicrobial Pathogenicity of Porphyromonas Gingivalis. Front. Oral Health 2024, Volume 5-2024. 10.3389/froh.2024.1404917

(7) Zhu, Y.; Dashper, S. G.; Chen, Y.-Y.; Crawford, S.; Slakeski, N.; Reynolds, E. C. Porphyromonas Gingivalis and Treponema Denticola Synergistic Polymicrobial Biofilm Development. PloS One 2013, 8 (8), e71727. 10.1371/journal.pone.0071727

(8) Kuboniwa, M.; Amano, A.; Hashino, E.; Yamamoto, Y.; Inaba, H.; Hamada, N.; Nakayama, K.; Tribble, G. D.; Lamont, R. J.; Shizukuishi, S. Distinct Roles of Long/Short Fimbriae and Gingipains in Homotypic Biofilm Development by Porphyromonas Gingivalis. BMC Microbiol. 2009, 9 (1), 105. 10.1186/1471-2180-9-105

(9) Park, Y.; Simionato, M. R.; Sekiya, K.; Murakami, Y.; James, D.; Chen, W.; Hackett, M.; Yoshimura, F.; Demuth, D. R.; Lamont, R. J. Short Fimbriae of Porphyromonas Gingivalis and Their Role in Coadhesion with Streptococcus Gordonii. Infect. Immun. 2005, 73 (7), 3983–3989. 10.1128/IAI.73.7.3983-3989.2005

(10) Hall, M.; Hasegawa, Y.; Yoshimura, F.; Persson, K. Structural and Functional Characterization of Shaft, Anchor, and Tip Proteins of the Mfa1 Fimbria from the Periodontal Pathogen Porphyromonas Gingivalis. Sci. Rep. 2018, 8 (1), 1793. 10.1038/s41598-018-20067-z

(11) Amano Atsuo; Nakamura Takayuki; Kimura Shigenobu; Morisaki Ichijiro; Nakagawa Ichiro; Kawabata Shigetada; Hamada Shigeyuki. Molecular Interactions of Porphyromonas gingivalisFimbriae with Host Proteins: Kinetic Analyses Based on Surface Plasmon Resonance. Infect. Immun. 1999, 67 (5), 2399–2405. 10.1128/iai.67.5.2399-2405.1999

(12) Maeda Kazuhiko; Nagata Hideki; Kuboniwa Masae; Kataoka Kosuke; Nishida Nobuko; Tanaka Muneo; Shizukuishi Satoshi. Characterization of Binding of Streptococcus Oralis Glyceraldehyde-3-Phosphate Dehydrogenase to Porphyromonas Gingivalis Major Fimbriae. Infect. Immun. 2004, 72 (9), 5475–5477. 10.1128/iai.72.9.5475-5477.2004

(13) Amano Atsuo; Shizukuishi Satoshi; Horie Hiroshi; Kimura Shigenobu; Morisaki Ichijiro; Hamada Shigeyuki. Binding of Porphyromonas gingivalisFimbriae to Proline-Rich Glycoproteins in Parotid Saliva via a Domain Shared by Major Salivary Components. Infect. Immun. 1998, 66 (5), 2072–2077. 10.1128/iai.66.5.2072-2077.1998

(14) Nakamura, T.; Amano, A.; Nakagawa, I.; Hamada, S. Specific Interactions between Porphyromonas Gingivalis Fimbriae and Human Extracellular Matrix Proteins. FEMS Microbiol. Lett. 1999, 175 (2), 267–272. 10.1016/S0378-1097(99)00205-0

(15) Hasegawa, Y.; Nagano, K. Porphyromonas Gingivalis FimA and Mfa1 Fimbriae: Current Insights on Localization, Function, Biogenesis, and Genotype. Jpn. Dent. Sci. Rev. 2021, 57, 190–200. 10.1016/j.jdsr.2021.09.003

(16) Hiramine, H.; Watanabe, K.; Hamada, N.; Umemoto, T. Porphyromonas Gingivalis 67-kDa Fimbriae Induced Cytokine Production and Osteoclast Differentiation Utilizing TLR2. FEMS Microbiol. Lett. 2003, 229 (1), 49–55. 10.1016/S0378-1097(03)00788-2

(17) Lin, X.; Wu, J.; Xie, H. *Porphyromonas Gingivalis* Minor Fimbriae Are Required for Cell-Cell Interactions. Infect. Immun. 2006, 74 (10), 6011–6015. 10.1128/iai.00797-06

(18) Lamont, R. J.; El-Sabaeny, A.; Park, Y.; Cook, G. S.; Costerton, J. W.; Demuth, D. R. Y. 2002. Role of the Streptococcus Gordonii SspB Protein in the Development of Porphyromonas Gingivalis Biofilms on Streptococcal Substrates. Microbiology, 148 (6), 1627–1636. 10.1099/00221287-148-6-1627

(19) Nagano, K.; Hasegawa, Y.; Murakami, Y.; Nishiyama, S.; Yoshimura, F. FimB Regulates FimA Fimbriation in Porphyromonas Gingivalis. J. Dent. Res. 2010, 89 (9), 903–908. 10.1177/0022034510370089

(20) Hasegawa, Y.; Iwami, J.; Sato, K.; Park, Y.; Nishikawa, K.; Atsumi, T.; Moriguchi, K.; Murakami, Y.; Lamont, R. J.; Nakamura, H.; Ohno, N.; Yoshimura, F. Anchoring and Length Regulation of Porphyromonas Gingivalis Mfa1 Fimbriae by the Downstream Gene Product Mfa2. Microbiology, 2009, 155, 3333–3347. 10.1099/mic.0.028928-0

(21) Nakagawa Ichiro; Amano Atsuo; Kuboniwa Masae; Nakamura Takayuki; Kawabata Shigetada; Hamada Shigeyuki. Functional Differences among FimA Variants of Porphyromonas Gingivalis and Their Effects on Adhesion to and Invasion of Human Epithelial Cells. Infect. Immun. 2002, 70 (1), 277–285. 10.1128/iai.70.1.277-285.2002

(22) Fujimoto, M.; Naiki, Y.; Sakae, K.; Iwase, T.; Miwa, N.; Nagano, K.; Nawa, H.; Hasegawa, Y. Structural and Antigenic Characterization of a Novel Genotype of Mfa1 Fimbriae in Porphyromonas Gingivalis. J. Oral Microbiol. 2023, 15.

(23) Xu, Q.; Shoji, M.; Shibata, S.; Naito, M.; Sato, K.; Elsliger, M.-A.; Grant, J. C.; Axelrod, H. L.; Chiu, H.-J.; Farr, C. L.; Jaroszewski, L.; Knuth, M. W.; Deacon, A. M.; Godzik, A.; Lesley, S. A.; Curtis, M. A.; Nakayama, K.; Wilson, I. A. A Distinct Type of Pilus from the Human Microbiome. Cell 2016, 165 (3), 690–703. 10.1016/j.cell.2016.03.016

(24) Shibata, S.; Matsunami, H.; Ouhara, K.; Taniguchi, Y.; Schreiber, M. T.; Villar-Brillones, A.; Nakayama, K.; Shoji, M.; Wolf, M. Cryo-EM Structure of the Native Assembled Mfa Type V Pilus from the Periodontal Pathogen Porphyromonas Gingivalis. Commun. Biol. 2026. 10.1038/s42003-026-10515-2

(25) Shibata, S.; Shoji, M.; Okada, K.; Matsunami, H.; Matthews, M. M.; Imada, K.; Nakayama, K.; Wolf, M. Structure of Polymerized Type V Pilin Reveals Assembly Mechanism Involving Protease-Mediated Strand Exchange. Nat. Microbiol. 2020, 5 (6), 830–837. 10.1038/s41564-020-0705-1

(26) Alaei, S. R.; Park, J. H.; Walker, S. G.; Thanassi, D. G. Peptide-Based Inhibitors of Fimbrial Biogenesis in Porphyromonas Gingivalis. Infect. Immun. 2019, 87 (3), 10.1128/iai.00750-18. 10.1128/iai.00750-18

(27) Lee, J. Y.; Miller, D. P.; Wu, L.; Casella, C. R.; Hasegawa, Y.; Lamont, R. J. Maturation of the Mfa1 Fimbriae in the Oral Pathogen Porphyromonas Gingivalis. Front. Cell. Infect. Microbiol. 2018, Volume 8-2018. 10.3389/fcimb.2018.00137

(28) Forero, M.; Yakovenko, O.; Sokurenko, E. V.; Thomas, W. E.; Vogel, V. Uncoiling Mechanics of Escherichia Coli Type I Fimbriae Are Optimized for Catch Bonds. PLOS Biol. 2006, 4 (9), e298. 10.1371/journal.pbio.0040298

(29) Krissinel, E.; Henrick, K. Inference of Macromolecular Assemblies from Crystalline State. J. Mol. Biol. 2007, 372 (3), 774–797. 10.1016/j.jmb.2007.05.022

(30) Heidler, T. V.; Ernits, K.; Ziolkowska, A.; Claesson, R.; Persson, K. Porphyromonas Gingivalis Fimbrial Protein Mfa5 Contains a von Willebrand Factor Domain and an Intramolecular Isopeptide. Commun. Biol. 2021, 4 (1), 106. 10.1038/s42003-020-01621-w

(31) Creasy, D. M.; Cottrell, J. S. Unimod: Protein Modifications for Mass Spectrometry. Proteomics 2004, 4 (6), 1534–1536. 10.1002/pmic.200300744

(32) Nagano, K.; Hasegawa, Y.; Yoshida, Y.; Yoshimura, F. A Major Fimbrilin Variant of Mfa1 Fimbriae in Porphyromonas Gingivalis. J. Dent. Res. 2015, 94 (8), 1143–1148. 10.1177/0022034515588275

(33) Gilchrist, C. L. M.; Mirdita, M.; Steinegger, M. Multiple Protein Structure Alignment at Scale with FoldMason. Science 2026, 391 (6784), 485–488. 10.1126/science.ads6733

(34) Gouet, P.; Courcelle, E.; Stuart, D. I.; Métoz, F. ESPript: Analysis of Multiple Sequence Alignments in PostScript. Bioinformatics 1999, 15 (4), 305–308. 10.1093/bioinformatics/15.4.305

(35) Robert, X.; Gouet, P. Deciphering Key Features in Protein Structures with the New ENDscript Server. Nucleic Acids Res. 2014, 42 (W1), W320–W324. 10.1093/nar/gku316

(36) Dhahri, S.; Ramonda, M.; Marlière, C. In-Situ Determination of the Mechanical Properties of Gliding or Non-Motile Bacteria by Atomic Force Microscopy under Physiological Conditions without Immobilization. PLOS ONE 2013, 8 (4), e61663. 10.1371/journal.pone.0061663

(37) Boudjemaa, R.; Steenkeste, K.; Canette, A.; Briandet, R.; Fontaine-Aupart, M.-P.; Marlière, C. Direct Observation of the Cell-Wall Remodeling in Adhering Staphylococcus Aureus 27217: An AFM Study Supported by SEM and TEM. Cell Surf. 2019, 5, 100018. 10.1016/j.tcsw.2019.100018

(38) Pakharukova, N.; Malmi, H.; Tuittila, M.; Dahlberg, T.; Ghosal, D.; Chang, Y.-W.; Myint, S. L.; Paavilainen, S.; Knight, S. D.; Lamminmäki, U.; Uhlin, B. E.; Andersson, M.; Jensen, G.; Zavialov, A. V. Archaic Chaperone–Usher Pili Self-Secrete into Superelastic Zigzag Springs. Nature 2022, 609 (7926), 335–340. 10.1038/s41586-022-05095-0

(39) Roky, M.; Trent, J. O.; Demuth, D. R. Identification of Functional Domains of the Minor Fimbrial Antigen Involved in the Interaction of Porphyromonas Gingivalis with Oral Streptococci. Mol. Oral Microbiol. 2020, 35 (2), 66–77. 10.1111/omi.12280

(40) Njoroge T; Genco R J; Sojar H T; Hamada N; Genco C A. A Role for Fimbriae in Porphyromonas Gingivalis Invasion of Oral Epithelial Cells. Infect. Immun. 1997, 65 (5), 1980–1984. 10.1128/iai.65.5.1980-1984.1997

(41) Malmi, H.; Pakharukova, N.; Paul, B.; Tuittila, M.; Ahmad, I.; Knight, S. D.; Uhlin, B. E.; Ghosal, D.; Zavialov, A. V. Antiparallel Stacking of Csu Pili Drives Acinetobacter Baumannii 3D Biofilm Assembly. Nat. Commun. 2026, 17 (1), 2508. 10.1038/s41467-026-68860-z

(42) Hasegawa, Y.; Nagano, K.; Murakami, Y.; Lamont, R. J. Purification of Native Mfa1 Fimbriae from Porphyromonas Gingivalis. In Periodontal Pathogens: Methods and Protocols; Nagano, K., Hasegawa, Y., Eds.; Springer US: New York, NY, 2021; pp 75–86. 10.1007/978-1-0716-0939-2_8

(43) Schorb, M.; Haberbosch, I.; Hagen, W. J. H.; Schwab, Y.; Mastronarde, D. N. Software Tools for Automated Transmission Electron Microscopy. Nat. Methods 2019, 16 (6), 471– 477. 10.1038/s41592-019-0396-9

(44) Punjani, A.; Rubinstein, J. L.; Fleet, D. J.; Brubaker, M. A. cryoSPARC: Algorithms for Rapid Unsupervised Cryo-EM Structure Determination. Nat. Methods 2017, 14 (3), 290– 296. 10.1038/nmeth.4169

(45) Meng, E. C.; Goddard, T. D.; Pettersen, E. F.; Couch, G. S.; Pearson, Z. J.; Morris, J. H.; Ferrin, T. E. UCSF ChimeraX: Tools for Structure Building and Analysis. Protein Sci. 2023, 32 (11), e4792. 10.1002/pro.4792

(46) Emsley, P.; Cowtan, K. It Coot: Model-Building Tools for Molecular Graphics. Acta Crystallogr. Sect. D 2004, 60 (12 Part 1), 2126–2132. 10.1107/S0907444904019158

(47) Afonine, P. V.; Poon, B. K.; Read, R. J.; Sobolev, O. V.; Terwilliger, T. C.; Urzhumtsev, A.; Adams, P. D. Real-Space Refinement in It PHENIX for Cryo-EM and Crystallography. Acta Crystallogr. Sect. D 2018, 74 (6), 531–544. 10.1107/S2059798318006551

(48) Williams, C. J.; Headd, J. J.; Moriarty, N. W.; Prisant, M. G.; Videau, L. L.; Deis, L. N.; Verma, V.; Keedy, D. A.; Hintze, B. J.; Chen, V. B.; Jain, S.; Lewis, S. M.; Arendall, W. B. 3rd; Snoeyink, J.; Adams, P. D.; Lovell, S. C.; Richardson, J. S.; Richardson, D. C. MolProbity: More and Better Reference Data for Improved All-Atom Structure Validation. Protein Sci. Publ. Protein Soc. 2018, 27 (1), 293–315. 10.1002/pro.3330

(49) Kim, H.-M.; Ranjit, D. K.; Walker, A. R.; Getachew, H.; Progulske-Fox, A.; Davey, M. E. A Novel Regulation of K-Antigen Capsule Synthesis in Porphyromonas Gingivalis Is Driven by the Response Regulator PG0720-Directed Antisense RNA. Front. Oral Health 2021, Volume 2-2021. 10.3389/froh.2021.701659

(50) Gardner, R. G.; Russell, J. B.; Wilson, D. B.; Wang, G. R.; Shoemaker, N. B. Use of a Modified Bacteroides-Prevotella Shuttle Vector to Transfer a Reconstructed Beta-1,4-D-Endoglucanase Gene into Bacteroides Uniformis and Prevotella Ruminicola B(1)4. Appl. Environ. Microbiol. 1996, 62 (1), 196–202. 10.1128/aem.62.1.196-202.1996

(51) Scott, J. C.; Klein, B. A.; Duran-Pinedo, A.; Hu, L.; Duncan, M. J. A Two-Component System Regulates Hemin Acquisition in Porphyromonas Gingivalis. PloS One 2013, 8 (9), e73351. 10.1371/journal.pone.0073351

(52) Abraham, M. J.; Murtola, T.; Schulz, R.; Páll, S.; Smith, J. C.; Hess, B.; Lindahl, E. GROMACS: High Performance Molecular Simulations through Multi-Level Parallelism from Laptops to Supercomputers. SoftwareX 2015, 1–2, 19–25. 10.1016/j.softx.2015.06.001

(53) Huang, J.; MacKerell Jr, A. D. CHARMM36 All-Atom Additive Protein Force Field: Validation Based on Comparison to NMR Data. J. Comput. Chem. 2013, 34 (25), 2135–2145. 10.1002/jcc.23354

(54) Jorgensen, W. L.; Chandrasekhar, J.; Madura, J. D.; Impey, R. W.; Klein, M. L. Comparison of Simple Potential Functions for Simulating Liquid Water. J. Chem. Phys. 1983, 79 (2), 926–935. 10.1063/1.445869

(55) Bussi, G.; Donadio, D.; Parrinello, M. Canonical Sampling through Velocity Rescaling. J. Chem. Phys. 2007, 126 (1), 014101. 10.1063/1.2408420

(56) Bernetti, M.; Bussi, G. Pressure Control Using Stochastic Cell Rescaling. J. Chem. Phys. 2020, 153 (11), 114107. 10.1063/5.0020514

(57) Michaud-Agrawal, N.; Denning, E. J.; Woolf, T. B.; Beckstein, O. MDAnalysis: A Toolkit for the Analysis of Molecular Dynamics Simulations. J. Comput. Chem. 2011, 32 (10), 2319–2327. 10.1002/jcc.21787

(58) Harris, C. R.; Millman, K. J.; van der Walt, S. J.; Gommers, R.; Virtanen, P.; Cournapeau, D.; Wieser, E.; Taylor, J.; Berg, S.; Smith, N. J.; Kern, R.; Picus, M.; Hoyer, S.; van Kerkwijk, M. H.; Brett, M.; Haldane, A.; del Río, J. F.; Wiebe, M.; Peterson, P.; Gérard-Marchant, P.; Sheppard, K.; Reddy, T.; Weckesser, W.; Abbasi, H.; Gohlke, C.; Oliphant, T. E. Array Programming with NumPy. Nature 2020, 585 (7825), 357–362. 10.1038/s41586-020-2649-2

(59) Saran, A.; Kim, H.-M.; Manning, I.; Hancock, M. A.; Schmitz, C.; Madej, M.; Potempa, J.; Sola, M.; Trempe, J.-F.; Zhu, Y.; Davey, M. E.; Zeytuni, N. Unveiling the Molecular Mechanisms of the Type IX Secretion System’s Response Regulator: Structural and Functional Insights. PNAS Nexus 2024, 3 (8), pgae316. 10.1093/pnasnexus/pgae316

(60) Al-Azzawi, Z. A. M.; Silver, N. R. G.; Mehra, S.; Niu, S.; Situ, C.; Luo, W.; Shlaifer, I.; Ingelsson, M.; Hyman, B. T.; Trempe, J.-F.; Durcan, T. M.; Watts, J. C.; Fon, E. A. α-Synuclein Purification Significantly Impacts Seed Amplification Assay Performance and Consistency. Acta Neuropathol. Commun. 2025, 13 (1), 225. 10.1186/s40478-025-02151-4

(61) Stevens, M. U.; Croteau, N.; Eldeeb, M. A.; Antico, O.; Zeng, Z. W.; Toth, R.; Durcan, T. M.; Springer, W.; Fon, E. A.; Muqit, M. M.; Trempe, J.-F. Structure-Based Design and Characterization of Parkin-Activating Mutations. Life Sci. Alliance 2023, 6 (6), e202201419. 10.26508/lsa.202201419

(62) Abramson, J.; Adler, J.; Dunger, J.; Evans, R.; Green, T.; Pritzel, A.; Ronneberger, O.; Willmore, L.; Ballard, A. J.; Bambrick, J.; Bodenstein, S. W.; Evans, D. A.; Hung, C.-C.; O’Neill, M.; Reiman, D.; Tunyasuvunakool, K.; Wu, Z.; Žemgulytė, A.; Arvaniti, E.; Beattie, C.; Bertolli, O.; Bridgland, A.; Cherepanov, A.; Congreve, M.; Cowen-Rivers, A. I.; Cowie, A.; Figurnov, M.; Fuchs, F. B.; Gladman, H.; Jain, R.; Khan, Y. A.; Low, C. M. R.; Perlin, K.; Potapenko, A.; Savy, P.; Singh, S.; Stecula, A.; Thillaisundaram, A.; Tong, C.; Yakneen, S.; Zhong, E. D.; Zielinski, M.; Žídek, A.; Bapst, V.; Kohli, P.; Jaderberg, M.; Hassabis, D.; Jumper, J. M. Accurate Structure Prediction of Biomolecular Interactions with AlphaFold 3. Nature 2024, 630 (8016), 493–500. 10.1038/s41586-024-07487-w

(63) Minh, B. Q.; Schmidt, H. A.; Chernomor, O.; Schrempf, D.; Woodhams, M. D.; von Haeseler, A.; Lanfear, R. IQ-TREE 2: New Models and Efficient Methods for Phylogenetic Inference in the Genomic Era. Mol. Biol. Evol. 2020, 37 (5), 1530–1534. 10.1093/molbev/msaa015

