## Supplementary information for "Structural tuning of native type V fimbriae shapes mechanical specialization in *Porphyromonas gingivalis*"

**Table S1. Cryo-EM data collection and refinement statistics.**

| | nFimA | nMfa1 | nMfa1$_{\Delta 51-64}$ |
|---|---|---|---|
| **PDB ID** | 37EC | 37EB | 37ED |
| **EMDB ID** | EMD-78110 | EMD-78109 | EMD-78111 |
| **Data collection and processing** | | | |
| Grids | C-flat R2/1 Cu-T | C-flat R2/1 Cu-T | C-flat R2/1 Cu-T |
| Vitrification method | FEI Vitrobot | FEI Vitrobot | FEI Vitrobot |
| Microscope | Titan Krios | Titan Krios | Glacios 2 |
| Magnification | 105,000x | 130,000x | 165,000x |
| Camera | Gatan K3 Camera | Gatan K3 Camera | Falcon 4i |
| Voltage (kV) | 300 | 300 | 200 |
| Electron exposure (e−/Å$^2$) | 60 | 60 | 61.88 |
| Number of frames | 40 | 40 | 40 |
| Defocus range (μm) | (-1.0) to (-2.5) | (-1.0) to (-2.5) | (-0.5) to (-2.5) |
| Pixel size (Å) | 0.855 | 0.675 | 0.720 |
| Number of micrographs | 9179 | 7020 | 6032 |
| Initial particle images (no.) | 4,329,302 | 1,150,574 | 504,616 |
| Symmetry | C1 | C1 | C1 |
| Final particle images (no.) | 4,043,200 | 505,844 | 53,358 |
| Map resolution (Å) | 2.92 | 2.90 | 3.78 |
| FSC threshold | 0.143 | 0.143 | 0.143 |
| Model resolution range (Å) (d FSC model 0/0.143/0.5) | 2.8/2.9/3.0 | 2.8/2.9/3.1 | 3.6/3.7/4.2 |
| **Refinement** | | | |
| Map resolution (Å) | 2.92 | 2.90 | 3.78 |
| FSC threshold | 0.143 | 0.143 | 0.143 |
| Map sharpening *B* factor (Å$^2$) | 146.9 | 108.0 | 70 |
| Non-hydrogen atoms | 7704 | 11793 | 11442 |
| Protein residues | 1,011 | 1539 | 1497 |
| ***B* factors (Å$^2$) (min/max/mean)** | | | |
| Protein | 3.29/151.93/39.84 | 87.80/266.21/156.63 | 93.09/252.31/178.28 |
| **r.m.s. deviations** | | | |
| Bond lengths (Å) | 0.003 | 0.003 | 0.003 |
| Bond angles (°) | 0.530 | 0.494 | 0.626 |

| | | | |
|---|---|---|---|
| **Model Validation** | | | |
| MolProbity score | 1.27 | 1.35 | 1.71 |
| Clashscore | 1.77 | 4.21 | 6.04 |
| Rotamer outliers (%) | 1.49 | 0.00 | 0.20 |
| **Ramachandran plot** | | | |
| Favored (%) | 96.71 | 97.58 | 96.78 |
| Allowed (%) | 3.29 | 2.42 | 3.02 |
| Disallowed (%) | 0 | 0 | 0 |

**Table S2. Primers used in this study.**

| Primers | Sequence (5' to 3') | Purpose |
|---|---|---|
| np-mfa1 fwd | TCGATAAGCTTGGATCCGCATGCCCCAGGGAAGCCGGTAAGACC | pT-COW-*np-mfa1* |
| np-mfa1 rev | GTTAGCGAGGTGCGGCCGGTCGACCCCTTAGAGATCAACCTCATAGGAATG | pT-COW-*np-mfa1* |
| np-mfa1$_{\Delta 51\text{-}64}$ fwd | GGGAAGTGCTCGTGCGGGAGAGTGGGCAGGAAAAG | Generating Mfa1 with G51-V64 deletion: pT-COW-*np-mfa1*$_{\Delta 51\text{-}64}$ |
| mfa1UP fwd | GACGTTGTAAAACGACGGCCAGTGCGCGTAAACTAATCGGTACCCTTTG | Generating *P. gingivalis* ATCC 33277 variant: $\Delta fimA::erm^R \Delta mfa1::cm^R$ |
| mfa1UP rev | GTGATTTTTTTCTCCATAAGCCAAATGTTTAAAAGGATTAATATTAAATTG | |
| mfa1DOWN fwd | GGGCGTAACATTCCTATGAGGTTGATCTCTAATTAGC | |
| mfa1DOWN rev | CCCGGGTACCGAGCTCGAATTCCAACGACATAAGTATCGTCAGGGC | |
| mfa1Cm fwd | CAATTTAATATTAATCCTTTTAAACATTTGGCTTATGGAGAAAAAAATCACTGGATATAC | |
| mfa1Cm rev | CTAATTAGAGATCAACCTCATAGGAATGTTACGCCCCGCCCTGCC | |
| pET28-mMfa1 fwd | GTGCCGCGCGGCAGCCATTGCGGGTGACGGACAGG | pET28a (+) -*mfa1* |
| pET28-mMfa1 rev | GTCGACGGAGCTCGAATTCGGATCCTTAGAGATCAACCTCATAGGAATG | pET28a (+) -*mfa1* |

**Table S3. Strains and plasmids used in this study**.

| **Strains (relevant genotype)** | **Sources** |
|---|---|
| ***E. coli strains*** | |
| TOP10 | NEB |
| BL21 | Life Technologies |
| S17-1 | [1] |
| ***P. gingivalis strains*** | |
| ATCC 33277 wild type | ATCC |
| ATCC 33277 *ΔfimA::erm*$^R$ | *fimA* gene is replaced by an erythromycin resistance (*erm*$^R$) cassette[2] |
| ATCC 33277 *Δmfa1::erm*$^R$ * | [3] |
| ATCC 33277 *ΔfimA::cm*$^R$*Δmfa1::erm*$^R$ ** | This study. Generated in *Δmfa1::erm*$^R$ [3] background by replacing *fimA* with a chloramphenicol-resistance cassette (*cm*$^R$). |
| ATCC 33277 *ΔfimA::erm*$^R$*Δmfa1::cm*$^R$ | This study. Generated in *ΔfimA::erm*$^R$ [2] background by replacing *mfa1* with a *cm*$^R$. Used for in trans complementation of the *mfa1* deletion with pT-COW carrying *mfa1*$_{Δ51\text{-}64}$ to assess assembly and properties of Mfa1$_{Δ51\text{-}64}$ fimbriae. |
| **Plasmids** | |
| pET28a+ (Kan$^R$) | Novagen |
| pUC19 (Amp$^R$) | NEB |
| pVA2198 (Erm$^R$ and Sp$^R$) | [4] |
| pT-COW (Amp$^R$ and Tc$^R$ in *E. coli*; Tc$^R$ in *P. gingivalis*; Mob+ Rep+) | [1] |
| pT-COW-*np*-*mfa1* | This study |
| pT-COW-*np*-*mfa1*$_{Δ51\text{-}64}$ | This study. Used for producing Mfa1$_{Δ51–64}$ fimbriae in *P. gingivalis.* |

* A ~6400 bp insertion downstream of *mfa1* disrupts expression of the downstream *mfa2*-*mfa5* genes[3].

** The parental *Δmfa1::erm*$^R$ strain carries the downstream insertion described above, so this strain also lacks functional expression of *mfa2-5*. Used for in trans complementation of the *mfa1* deletion with pT-COW carrying wild-type *mfa1,* resulting in assembly of elongated Mfa1 fimbriae in the absence of a functional Mfa2.

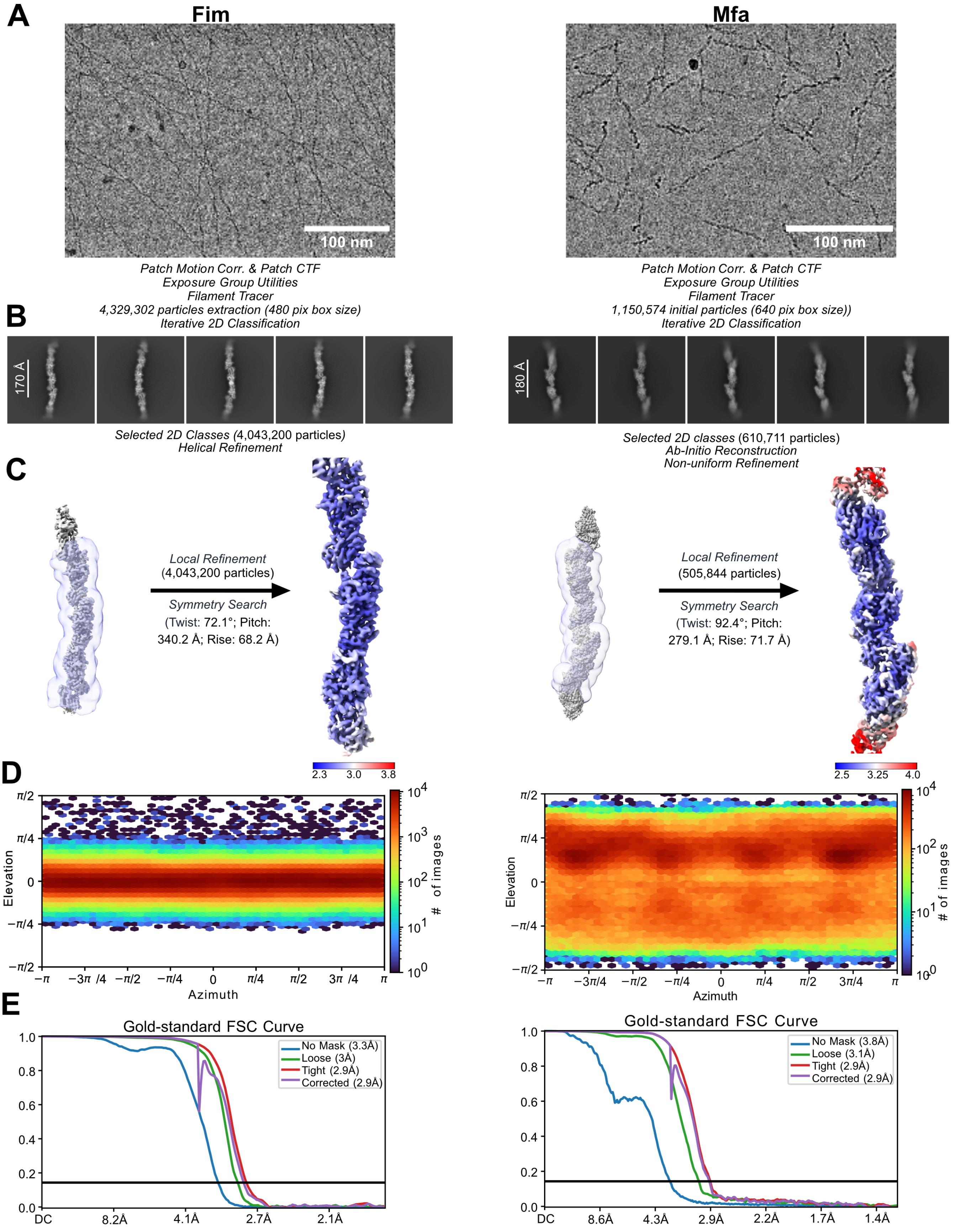
A
Fim
Mfa
100 nm
100 nm
Patch Motion Corr. & Patch CTF
Exposure Group Utilities
Filament Tracer
4,329,302 particles extraction (480 pix box size)
Iterative 2D Classification
Patch Motion Corr. & Patch CTF
Exposure Group Utilities
Filament Tracer
1,150,574 initial particles (640 pix box size))
Iterative 2D Classification
B
170 Å
180 Å
Selected 2D Classes (4,043,200 particles)
Helical Refinement
Selected 2D classes (610,711 particles)
Ab-Initio Reconstruction
Non-uniform Refinement
C
Local Refinement
(4,043,200 particles)
Symmetry Search
(Twist: 72.1°; Pitch: 340.2 Å; Rise: 68.2 Å)
Local Refinement
(505,844 particles)
Symmetry Search
(Twist: 92.4°; Pitch: 279.1 Å; Rise: 71.7 Å)
2.3 3.0 3.8
2.5 3.25 4.0
D
Elevation
Azimuth
### of images
E
Gold-standard FSC Curve
No Mask (3.3Å)
Loose (3Å)
Tight (2.9Å)
Corrected (2.9Å)
DC 8.2Å 4.1Å 2.7Å 2.1Å
Gold-standard FSC Curve
No Mask (3.8Å)
Loose (3.1Å)
Tight (2.9Å)
Corrected (2.9Å)
DC 8.6Å 4.3Å 2.9Å 2.2Å 1.7Å 1.4Å

**Figure S1**. **Cryo-EM data processing and refinement workflows for native Fim and Mfa fimbriae**. Fim and Mfa are shown on the left and right, respectively. (**A**) Representative cryo-EM micrographs; scale bars, 100 nm. (**B**) Representative 2D class averages. For Fim, 4,329,302 particles were extracted using a 480-pixel box, and 4,043,200 particles were retained after 2D classification. For Mfa, 1,150,574 particles were extracted using a 640-pixel box, and 610,711 particles were retained. (**C**) 3D reconstruction and refinement. Consensus maps are shown with the masks used for subsequent local refinement displayed as semitransparent violet surfaces. Local refinement of 4,043,200 Fim particles and 505,844 Mfa particles produced the final maps. Symmetry searching yielded a helical twist of 72.1°, a rise of 68.2 Å, and a pitch of 340.2 Å for Fim, and a twist of 92.4°, a rise of 71.7 Å, and a pitch of 279.1 Å for Mfa. (**D**) Viewing-direction distributions for particles contributing to the final refinements. (**E**) Gold-standard Fourier shell correlation curves; the horizontal line indicates the FSC = 0.143 criterion.

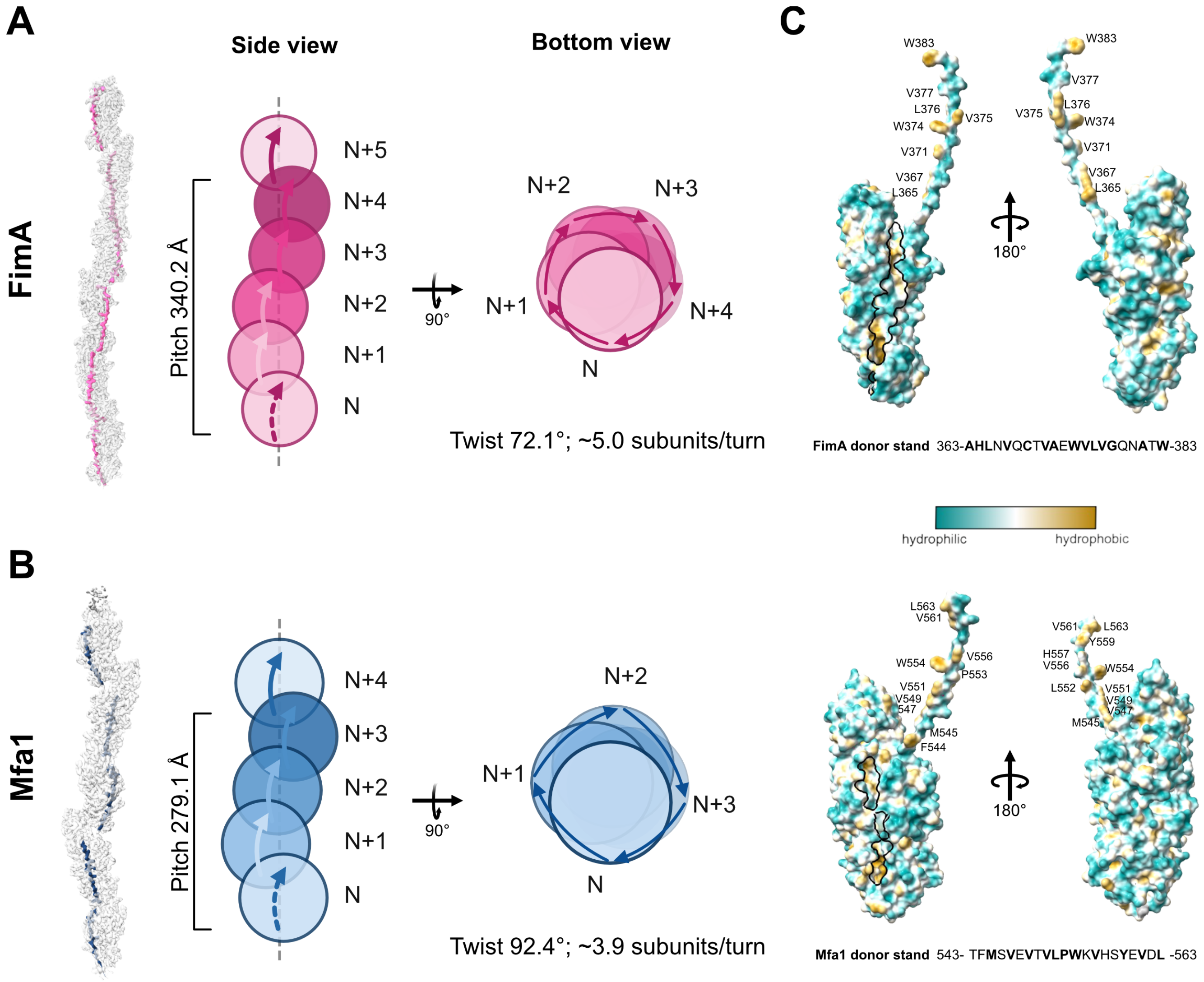
A
Side view
Bottom view
FimA
Pitch 340.2 Å
N+5
N+4
N+3
N+2
N+1
N
90°
Twist 72.1°; ~5.0 subunits/turn
C
W383
V377
L376
W374
V375
V371
V367
L365
180°
FimA donor stand 363-AHLNVQCTVAEWVLVGQNATW-383
hydrophilic
hydrophobic
B
Mfa1
Pitch 279.1 Å
Twist 92.4°; ~3.9 subunits/turn
L563
V561
Y559
H557
V556
W554
P553
L552
V551
V549
547
M545
F544
Mfa1 donor stand 543- TFMSVEVTVLPWKVHSYEVDL -563

**Figure S2. Comparison of stalk subunit packing in Fim and Mfa fimbriae**. Side and bottom views illustrate the helical packing of consecutive subunits in Fim (A) and Mfa (B) fimbriae, respectively. Filament models are shown on the left, with simplified packing schematics shown in the center and right. Composite cryo-EM volumes were generated in ChimeraX[5] using the volume maximum function to merge two maps spanning three consecutive subunits. Donor strands of FimA and Mfa1 are colored pink and blue on the map. Individual subunits are labeled sequentially from N to N+i, and curved arrows indicate the direction of helical progression and subunit rotation around the filament axis. In the side views, the grey dashed line marks the central filament axis, and the bracketed measurements indicate the helical pitch. Bottom-view projections show the angular positions occupied by successive subunits around the filament circumference. FimA has a pitch of 340.2 Å, a rise of 68.2 Å, a twist of 72.1°, and ~5.0 subunits per turn, whereas Mfa1 has a pitch of 279.1 Å, a rise of 71.7 Å, a twist of 92.4°, and ~3.9 subunits per turn. (**C**) Surface representations of polymerized FimA (**top**), and Mfa1 (**bottom**) stalk pilin subunits colored by hydrophobicity, with hydrophilic regions in cyan, hydrophobic regions in goldenrod, and intermediate regions in white. For each subunit, two views related by a 180° rotation are shown for each protein. Black outlines mark the C-terminal donor-strand binding groove, and selected donor-strand residues are labeled.

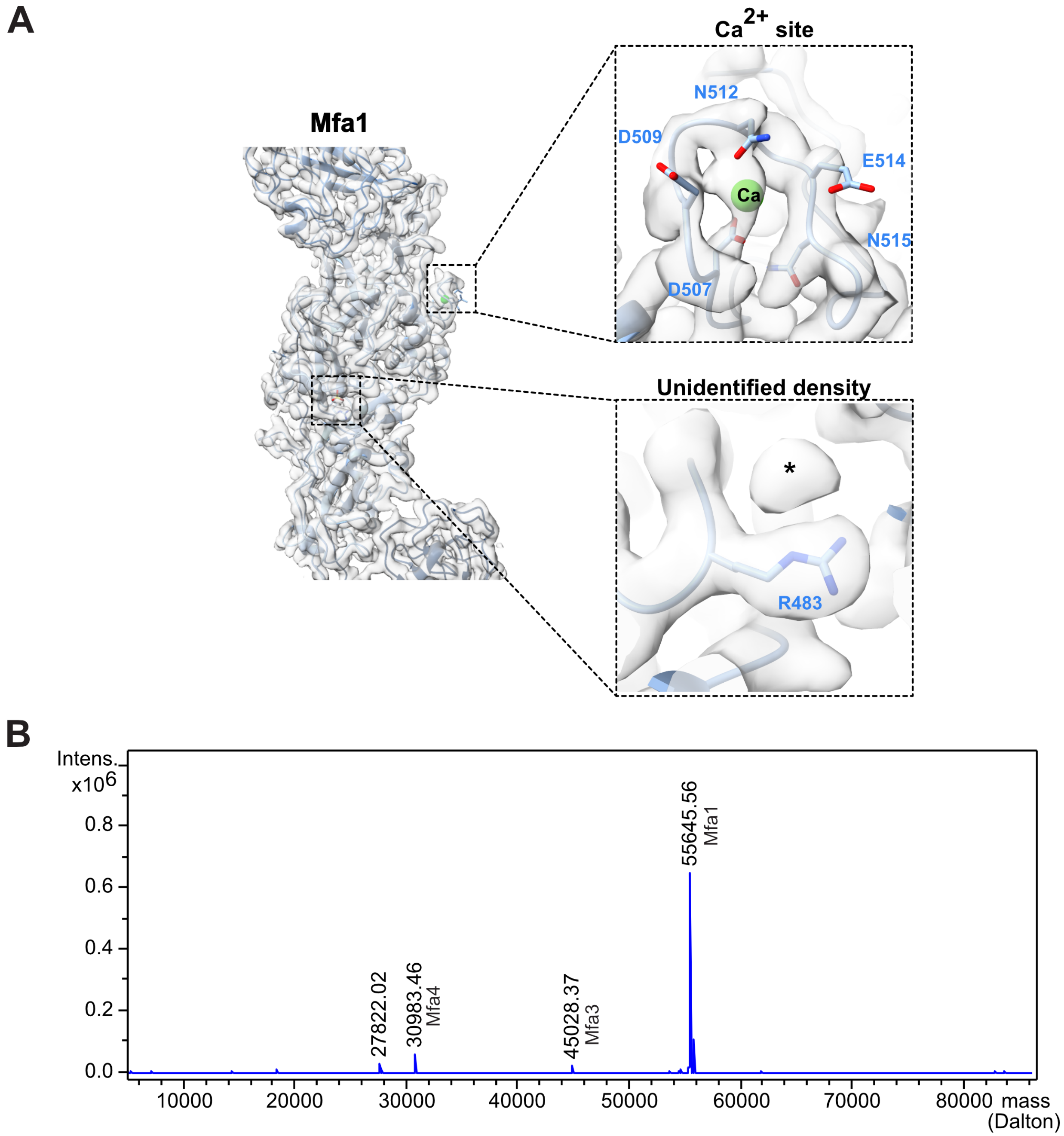

A
Mfa1
Ca2+ site
N512
D509
E514
Ca
N515
D507
Unidentified density
*
R483
B
Intens.
x10^6
0.8
0.6
0.4
0.2
0.0
55645.56
Mfa1
27822.02
30983.46
Mfa4
45028.37
Mfa3
10000
20000
30000
40000
50000
60000
70000
80000
mass
(Dalton)

**Figure S3. Local cryo-EM density and non-protein density features of the native Mfa fimbriae filament.** Overall view of a single Mfa1 subunit within the native Mfa filament, with enlarged views of two non-protein density features. The upper panel shows an assigned $Ca^{2+}$-binding site, with the calcium ion shown as a green sphere and coordinating residues as sticks. The lower inset shows an unidentified density (*) adjacent to R483. This density lies in a pocket corresponding to a site modeled as acetate-bound in the monomeric Mfa1 crystal structure (PDB 5NF2) but was not assigned to a specific ligand in the native Mfa1 cryo-EM model. Cryo-EM density is displayed as a semitransparent gray surface. (**B**) Representative deconvolution for 1 µg purified nMfa analyzed by intact LC-MS. The predominant peak detected corresponded to the mature Mfa1 protein species (55,647.3 Da theoretical). Minor species with observed masses of 30,983.5 and 45,028.4 Da are consistent with Mfa4 and Mfa3 (theoretical 30983.89 Da and 45029.54 Da), respectively. The peak at mass 27,822.02 Da corresponds to the doubly charged Mfa1 ion, $[M + 2H]^{2+}$.

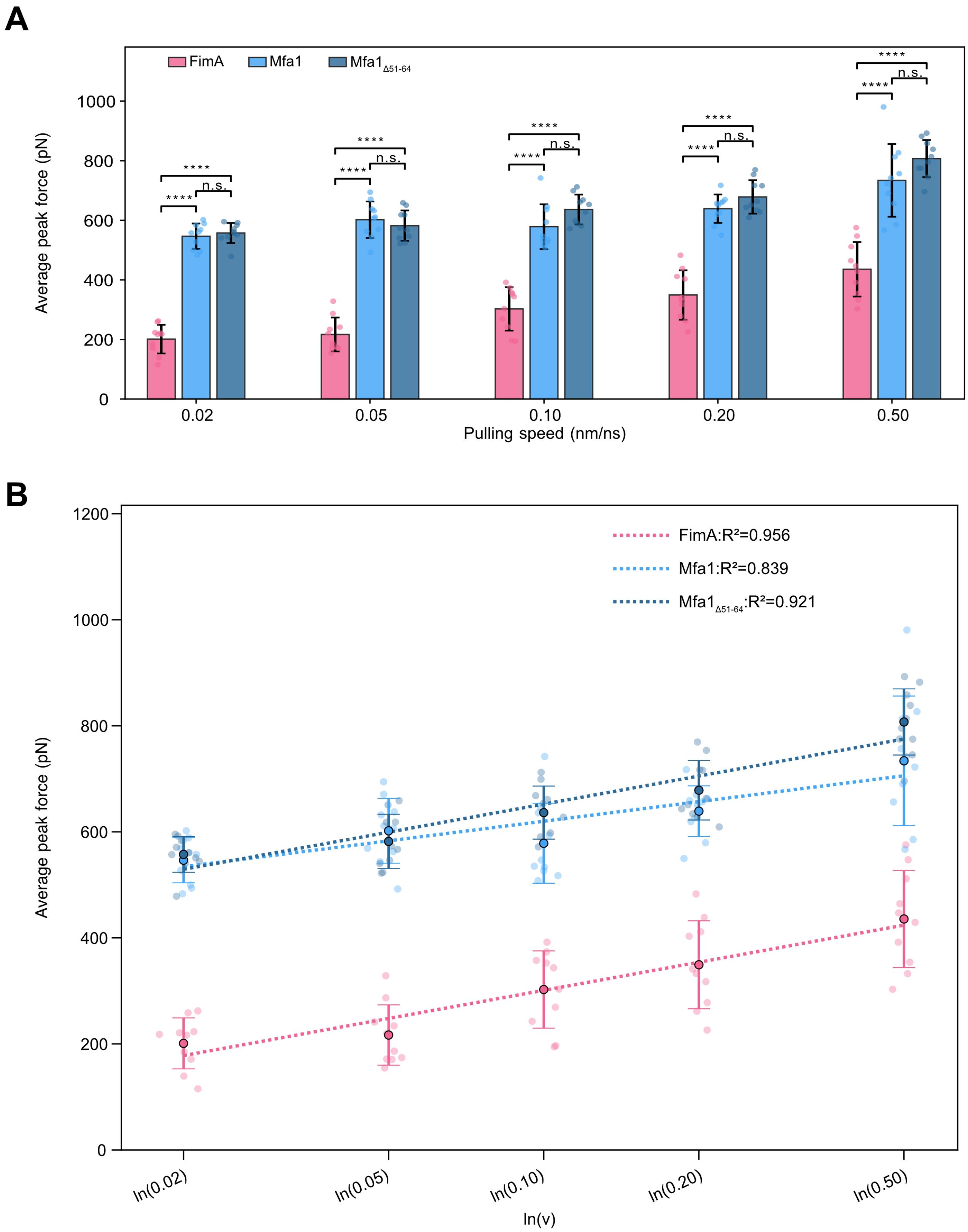
A
FimA
Mfa1
Mfa1Δ51-64
Average peak force (pN)
Pulling speed (nm/ns)
0.02
0.05
0.10
0.20
0.50
****
n.s.
B
FimA:R²=0.956
Mfa1:R²=0.839
Mfa1Δ51-64:R²=0.921
Average peak force (pN)
ln(0.02)
ln(0.05)
ln(0.10)
ln(0.20)
ln(0.50)
ln(v)

**Figure S4. Initial peak force of FimA, Mfa1, and Mfa1$_{\Delta 51\text{-}64}$ stalk fragments across pulling speeds. (A)** Initial peak force measured in SMD pulling simulations at speeds of 0.02, 0.05, 0.10, 0.20, and 0.50 nm ns$^{-1}$. The initial peak force over the analyzed extension range was extracted from each replicate. Bars indicate the mean of per-replicate initial peak forces for each speed (n=10 per speed), with individual simulation replicates shown as overlaid circles. Error bars denote the sample standard deviation (SD). Statistical significance was assessed by pairwise Welch's t-tests within each pulling speed with Holm correction, ****$P < 0.0001$, n.s., not significant. (**B**) Bell-Evans-type[6] analysis showing the relationship between the mean per-replicate initial peak force and the natural logarithm of pulling speed, where v is expressed in nm ns$^{-1}$. Large, outlined circles show group means, smaller transparent circles represent individual forces, and error bars indicate SD. Dotted lines show unweighted least-squares fits of mean maximum force versus ln(v). The fitted relationships were: FimA, $F = 76.4 \ln(v) + 477.1$, $R^2 = 0.96$; Mfa1, $F = 53.3 \ln(v) + 742.8$, $R^2 = 0.84$; and Mfa1$_{\Delta 51\text{–}64}$, $F = 76.4 \ln(v) + 828.1$, $R^2 = 0.92$.

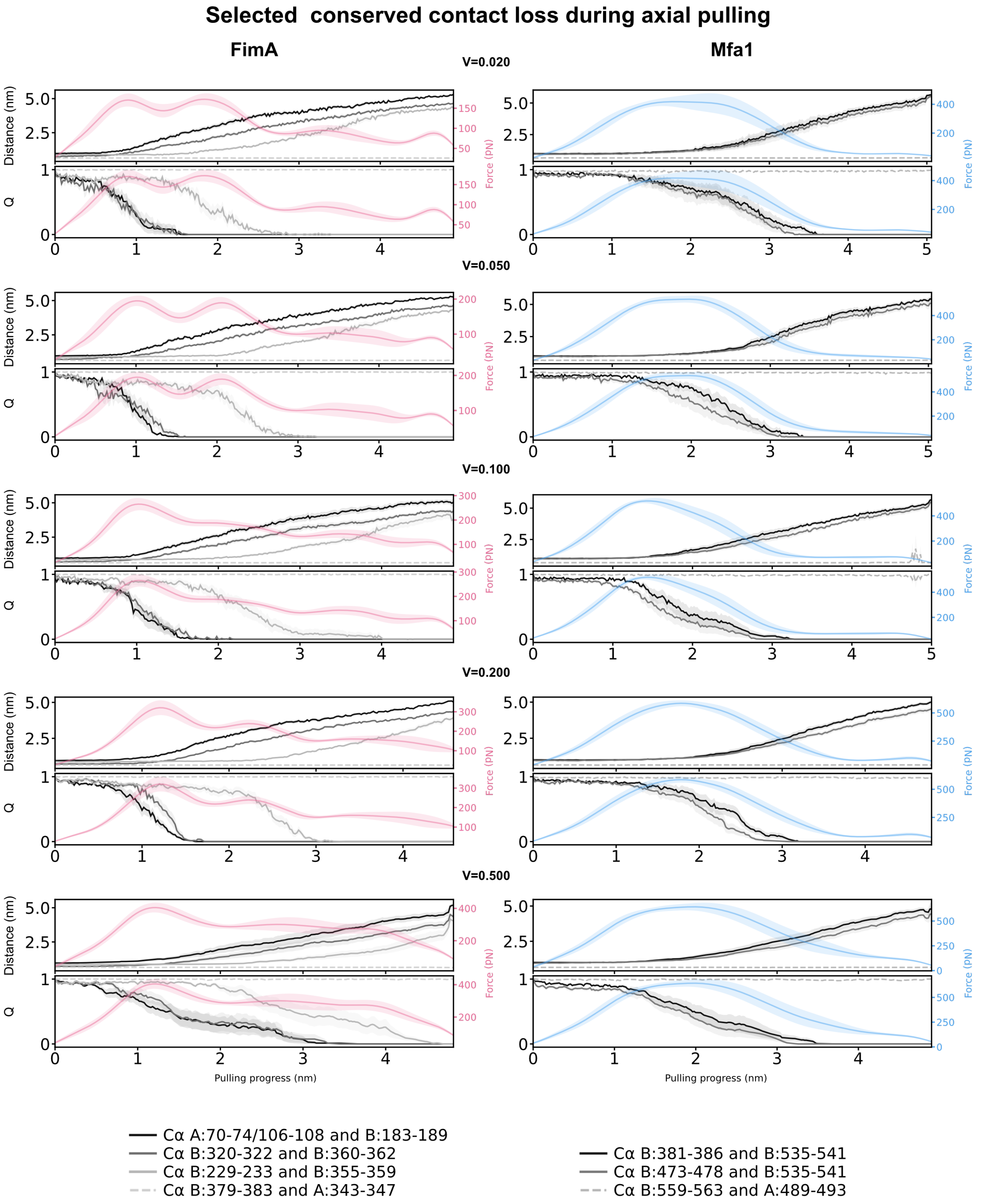

Selected conserved contact loss during axial pulling
FimA
Mfa1
V=0.020
V=0.050
V=0.100
V=0.200
V=0.500
Distance (nm)
Q
Force (PN)
Pulling progress (nm)
Cα A:70-74/106-108 and B:183-189
Cα B:320-322 and B:360-362
Cα B:229-233 and B:355-359
Cα B:379-383 and A:343-347
Cα B:381-386 and B:535-541
Cα B:473-478 and B:535-541
Cα B:559-563 and A:489-493

**Figure S5. Pulling-rate dependence of selected Cα-contact dynamics in FimA and Mfa1 stalk fragments.** Selected Cα-Cα contact pairs were monitored during SMD pulling of FimA and Mfa1 two-subunit stalk fragments at pulling speeds of 0.020, 0.050, 0.100, 0.200, and 0.50 nm/ns. For each pulling speed, the upper panels show the change in Cα-Cα distance for residue pairs as a function of pulling progress, with the corresponding force-extension profile overlaid on the right y-axis. The lower panels show the normalized Cα-contact Q score for the same residue pairs, where values near 1 indicate retention of the native contact and values approaching 0 indicate contact separation. FimA force profiles are shown in pink and Mfa1 force profiles in blue. Black and gray traces correspond to the residue pairs listed below each column. Lines represent means across ten independent trajectories, and shaded regions indicate standard error of the mean (SEM). Dashed light-gray traces denote the C-terminal donor-strand associated residue pairs, which remained stable during pulling.

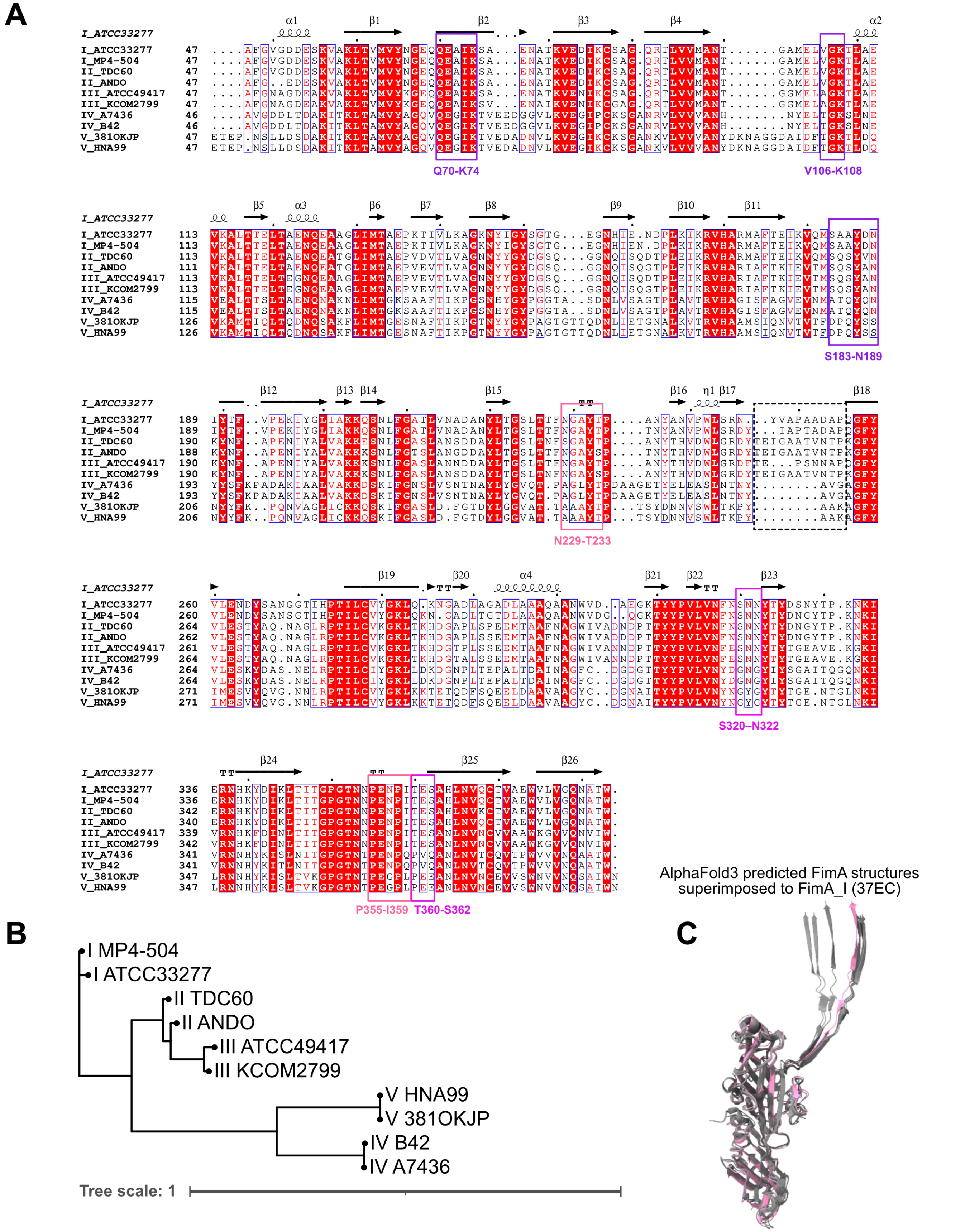

A
I_ATCC33277
I_MP4-504
II_TDC60
II_ANDO
III_ATCC49417
III_KCOM2799
IV_A7436
IV_B42
V_381OKJP
V_HNA99
Q70-K74
V106-K108
S183-N189
N229-T233
S320–N322
P355-I359
T360-S362
B
I MP4-504
I ATCC33277
II TDC60
II ANDO
III ATCC49417
III KCOM2799
V HNA99
V 381OKJP
IV B42
IV A7436
Tree scale: 1
C
AlphaFold3 predicted FimA structures
superimposed to FimA_I (37EC)

**Figure S6. Sequence conservation of mechanically responsive contact regions in mature FimA.** (**A**) Structure-guided multiple-sequence alignment of representative FimA proteins from types I-V genotypes: type I, ATCC 33277 and MP4-504 strains; type II, TDC60 and Ando strains; type III, ATCC 49417 and KCOM2799 strains; type IV, B42 and A7436 strains; and type V, HNA99 and 381OKJP strains. Secondary-structure elements and residue numbering refer to FimA from ATCC 33277. Purple-outlined boxes identify the Q70-K74, V106-K108, and S183-N189 contact regions. Magenta-outlined boxes identify the S320-N322/T360-S362 intra-subunit network, which is also disrupted during the early response, and the N229-T233/P355-I359 network, which is disrupted later during extension. The Q70-K74/V106-K108/S183-N189 and N229-T233/P355-I359 networks contain highly conserved residues across the FimA types examined, whereas the S320-N322/T360-S362 network contains conserved positions interspersed with type-associated substitutions. The structure-guided alignment was generated using FoldMason[7] and visualized with ESPript 3.0[8]; conserved and similar residues are highlighted according to ESPript conventions. (**B**) The phylogenetic tree summarizes relationships among the ten FimA variants. (**C**) Structural superposition of FimA proteins in polymerized form. The structural overlay yielded an alignment-mapped Cα RMSD of 1.26-1.70 Å relative to FimA-I from ATCC 33277.

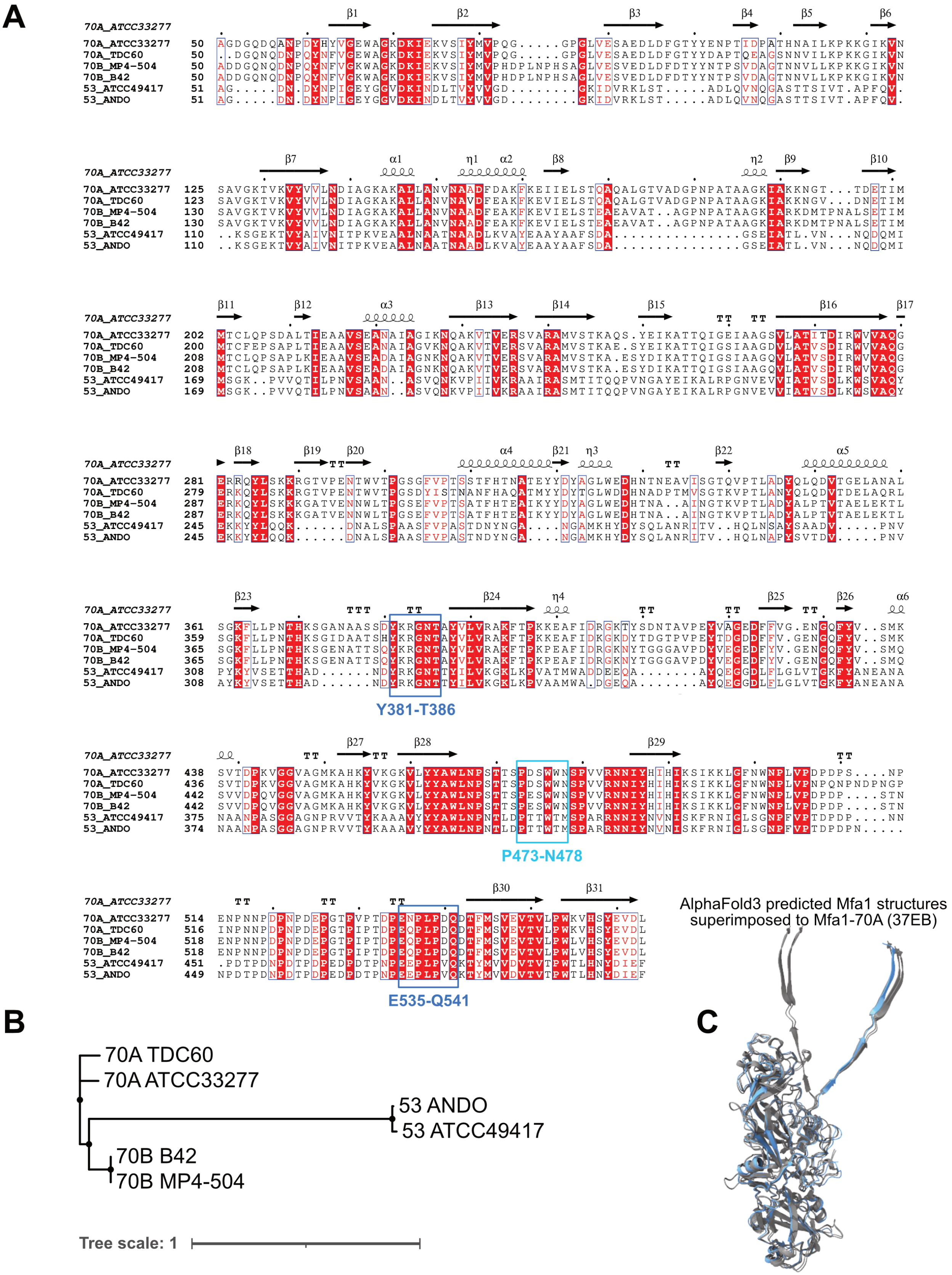
A
B
C
70A_ATCC33277
70A_TDC60
70B_MP4-504
70B_B42
53_ATCC49417
53_ANDO
Y381-T386
P473-N478
E535-Q541
AlphaFold3 predicted Mfa1 structures superimposed to Mfa1-70A (37EB)
70A TDC60
70A ATCC33277
53 ANDO
53 ATCC49417
70B B42
70B MP4-504
Tree scale: 1

**Figure S7. Sequence conservation of mechanically responsive contact regions in mature Mfa1.** (**A**) Structure-guided multiple-sequence alignment of representative Mfa1-70A proteins from ATCC 33277 and TDC60 strains, Mfa1-70B proteins from MP4-504 and B42 strains, and Mfa1-53 proteins from ATCC 49417 and Ando strains. Secondary-structure elements and residue numbering refer to Mfa1-70A from ATCC 33277. Colored boxes identify the mechanically responsive segments monitored in the SMD contact-loss analysis. Blue-outlined boxes mark Y381-T386 and E535-Q541, which form the contact pair associated with the major contact-loss transition; the light-cyan-outlined box marks P473–N478, an additional monitored interaction with E535-Q541. Y381-T386 and E535-Q541 are highly conserved among the Mfa1 variants examined, whereas P473-N478 is broadly conserved but contains limited group-associated substitutions. Box colors correspond to the respective contact-loss traces in the SMD analysis. The structure-guided alignment was generated using FoldMason[7] and visualized with ESPript 3.0[8]; conserved and similar residues are highlighted according to ESPript conventions. (**B**) The phylogenetic tree summarizes relationships among the six Mfa1 variants. (**C**) Structural superposition of Mfa1 proteins in polymerized form. The structural overlay yielded an alignment-mapped Cα RMSD of 1.18 to 1.92 Å relative to Mfa1-70A from ATCC 33277.

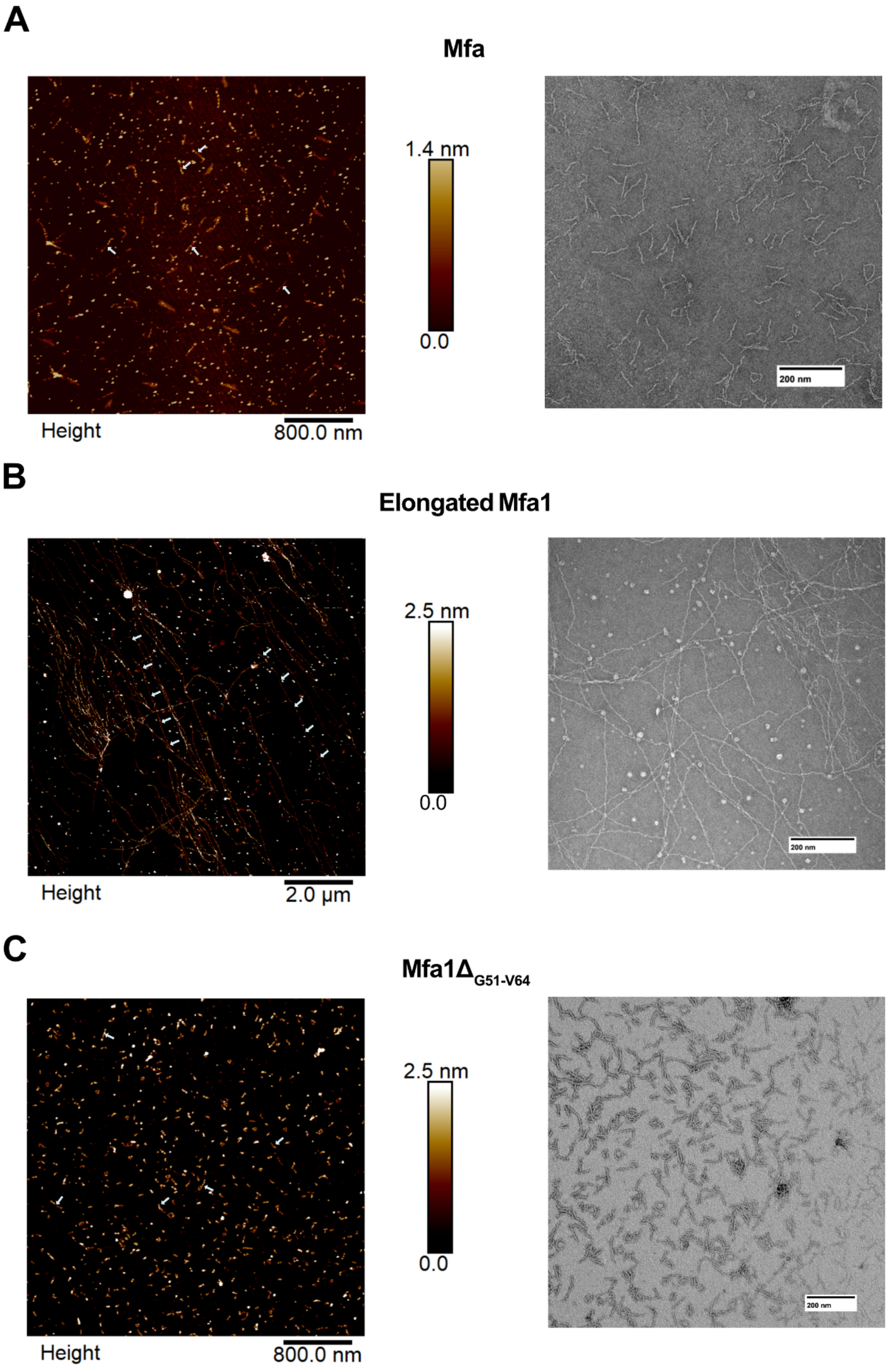
A
Mfa
1.4 nm
0.0
Height
800.0 nm
200 nm
B
Elongated Mfa1
2.5 nm
0.0
Height
2.0 µm
200 nm
C
Mfa1Δ$_{G51-V64}$
2.5 nm
0.0
Height
800.0 nm
200 nm

**Figure S8. (A-C)** Representative atomic force microscopy (AFM) height images (*left*) and negative-stain transmission electron microscopy (TEM) images (*right*) of purified native Mfa, elongated Mfa1, and Mfa1$_{\Delta 51–64}$ fimbriae, respectively. Multiple arrows trace the path of selected elongated Mfa filaments, whereas individual arrows mark representative shorter Mfa and Mfa1$_{\Delta 51-64}$ filaments. Scale bars are indicated in each panel.

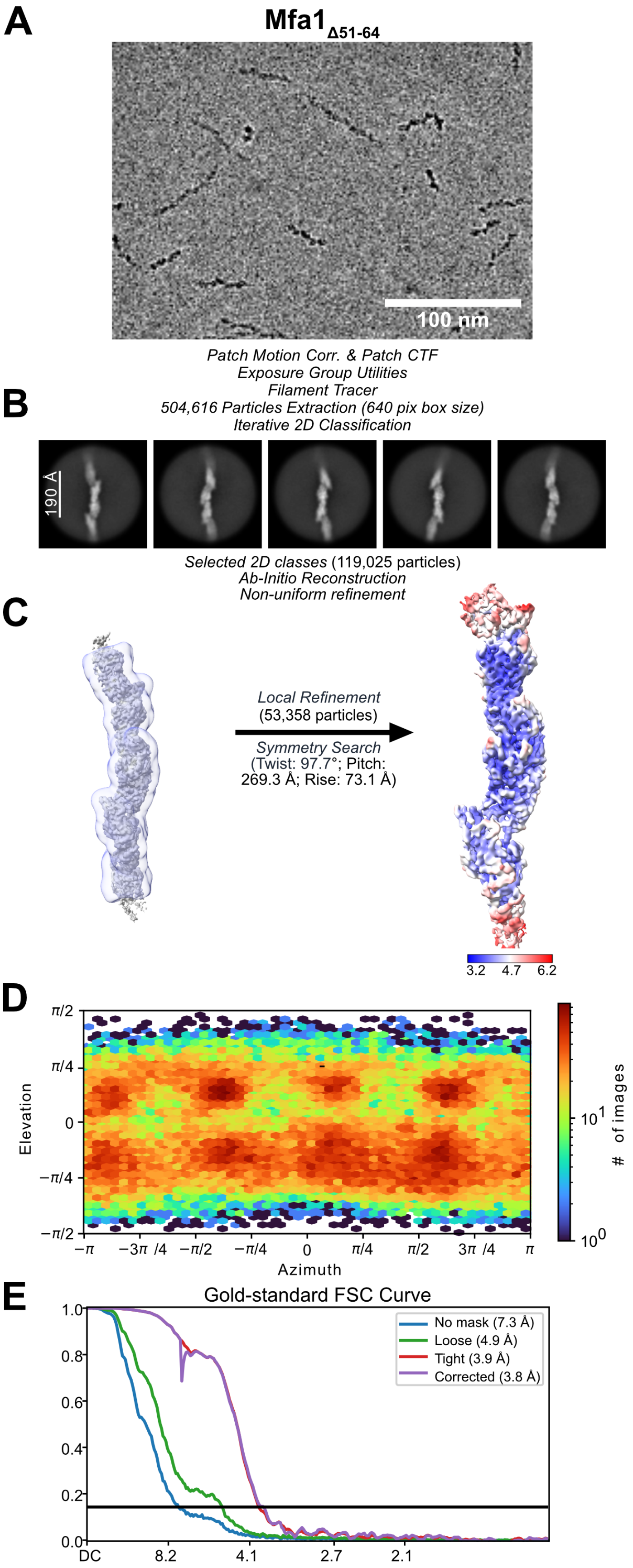
A
Mfa1Δ51-64
100 nm
Patch Motion Corr. & Patch CTF
Exposure Group Utilities
Filament Tracer
504,616 Particles Extraction (640 pix box size)
Iterative 2D Classification
B
190 Å
Selected 2D classes (119,025 particles)
Ab-Initio Reconstruction
Non-uniform refinement
C
Local Refinement
(53,358 particles)
Symmetry Search
(Twist: 97.7°; Pitch: 269.3 Å; Rise: 73.1 Å)
3.2
4.7
6.2
D
Elevation
Azimuth
### of images
E
Gold-standard FSC Curve
No mask (7.3 Å)
Loose (4.9 Å)
Tight (3.9 Å)
Corrected (3.8 Å)
DC
8.2
4.1
2.7
2.1

**Figure S9. Cryo-EM data processing and refinement workflow for Mfa1$_{\Delta 51\text{-}64}$ fimbriae. (A)** Representative cryo-EM micrograph of Mfa1$_{\Delta 51\text{-}64}$ filaments; scale bar, 100 nm. **(B)** Representative 2D class averages. A total of 504,616 particles were extracted using a 640-pixel box, of which 119,025 particles were retained after 2D classification. **(C)** 3D reconstruction and refinement. The consensus map obtained by ab initio reconstruction and nonuniform refinement is shown with the mask used for subsequent local refinement displayed as a semitransparent violet surface. Local refinement of 53,358 particles produced the final map, which is colored according to local resolution. Symmetry searching yielded a pitch of 269.3 Å, a rise of 73.1 Å, a helical twist of 97.7°, and ~3.7 subunits per turn. **(D)** Viewing-direction distribution for particles contributing to the final refinement. **(E)** Gold-standard Fourier shell correlation curves; the horizontal line indicates the FSC = 0.143 criterion.

#### References

(1) Matsumoto-Mashimo, C.; Guerout, A.-M.; Mazel, D. A New Family of Conditional Replicating Plasmids and Their Cognate Escherichia Coli Host Strains. *Res Microbiol* **2004**, *155* (6), 455–461. https://doi.org/10.1016/j.resmic.2004.03.001

(2) Malek R; Fisher J G; Caleca A; Stinson M; van Oss C J; Lee J Y; Cho M I; Genco R J; Evans R T; Dyer D W. Inactivation of the Porphyromonas Gingivalis fimA Gene Blocks Periodontal Damage in Gnotobiotic Rats. *Journal of Bacteriology* **1994**, *176* (4), 1052–1059. https://doi.org/10.1128/jb.176.4.1052-1059.1994

(3) Lamont, R. J.; El-Sabaeny, A.; Park, Y.; Cook, G. S.; Costerton, J. W.; Demuth, D. R. Y. 2002. Role of the Streptococcus Gordonii SspB Protein in the Development of Porphyromonas Gingivalis Biofilms on Streptococcal Substrates. *Microbiology, 148* (6), 1627–1636. https://doi.org/10.1099/00221287-148-6-1627

(4) Fletcher H M; Schenkein H A; Morgan R M; Bailey K A; Berry C R; Macrina F L. Virulence of a Porphyromonas Gingivalis W83 Mutant Defective in the prtH Gene. *Infection and Immunity* **1995**, *63* (4), 1521–1528. https://doi.org/10.1128/iai.63.4.1521-1528.1995

(5) Meng, E. C.; Goddard, T. D.; Pettersen, E. F.; Couch, G. S.; Pearson, Z. J.; Morris, J. H.; Ferrin, T. E. UCSF ChimeraX: Tools for Structure Building and Analysis. *Protein Science* **2023**, *32* (11), e4792. https://doi.org/10.1002/pro.4792

(6) Akhunzada, M. J.; Yoon, H. J.; Deb, I.; Braka, A.; Wu, S. Bell-Evans Model and Steered Molecular Dynamics in Uncovering the Dissociation Kinetics of Ligands Targeting G-Protein-Coupled Receptors. *Scientific Reports* **2022**, *12* (1), 15972. https://doi.org/10.1038/s41598-022-20065-2

(7) Gilchrist, C. L. M.; Mirdita, M.; Steinegger, M. Multiple Protein Structure Alignment at Scale with FoldMason. *Science* **2026**, *391* (6784), 485–488. https://doi.org/10.1126/science.ads6733

(8) Gouet, P.; Courcelle, E.; Stuart, D. I.; Métoz, F. ESPript: Analysis of Multiple Sequence Alignments in PostScript. *Bioinformatics* **1999**, *15* (4), 305–308. https://doi.org/10.1093/bioinformatics/15.4.305
